# Predictive features of gene expression variation reveal a mechanistic link between expression variation and differential expression

**DOI:** 10.1101/2020.02.10.942276

**Authors:** Olga M. Sigalova, Amirreza Shaeiri, Mattia Forneris, Eileen E.M. Furlong, Judith B Zaugg

## Abstract

For most biological processes, organisms must respond to extrinsic cues, while maintaining essential gene expression programs. Although studied extensively in single cells, it is still unclear how variation is controlled in multicellular organisms. Here, we used a machine-learning approach to identify genomic features that are predictive of genes with high versus low variation in their expression across individuals, using bulk data to remove stochastic cell-to-cell variation. Using embryonic gene expression across 75 *Drosophila* isogenic lines, we identify features predictive of expression variation, while controlling for expression level. Genes with low variation fall into two classes, indicating they employ different mechanisms to maintain a robust expression. In contrast, genes with high variation seem to lack both types of stabilizing mechanisms. Applying the framework to human tissues from GTEx revealed similar predictive features, indicating that promoter architecture is an ancient mechanism to control expression variation. Remarkably, expression variation features could also predict differential expression upon stress in both *Drosophila* and human. Differential gene expression signatures may therefore be partially explained by genetically encoded gene-specific features, unrelated to the studied treatment.

## Introduction

Living systems have a remarkable capacity to give rise to robust and highly reproducible phenotypes. Perhaps the most striking example of this is the process of embryogenesis, where fertilized eggs give rise to stereotypic body plans despite segregating genetic variants and moderate differences in environmental conditions (e.g. water temperature for fish, mothers diet for humans). This phenomenon led Waddington to propose that developmental reactions are canalized, which buffers them to withstand such variation without alterations in embryonic development (Waddington 1942). In agreement with this, variation in gene expression is an evolvable trait under selection pressure (Lehner 2008; Fraser et al. 2004; Metzger et al. 2015).

Gene expression variation can arise from a multitude of stochastic, environmental and genetic factors (Eling, Morgan, and Marioni 2019; Raser and O’Shea 2005; Félix and Barkoulas 2015; S. Huang 2009). For some genes, expression variation is tolerated, without obvious effects on fitness, or can even be beneficial, for example in stress response or for stochastic cell fate decisions (Macneil and Walhout 2011; Raj and van Oudenaarden 2008; Blake et al. 2006). In other cases, variation in gene expression is detrimental and must be tightly regulated, for example for essential genes (Fraser et al. 2004) and genes that reduce fitness in heterozygous mutants (Batada and Hurst 2007). This suggests that there are inherent mechanisms that modulate variation in gene expression, either attenuating or amplifying it (Fig 1a).

**Figure 1.**
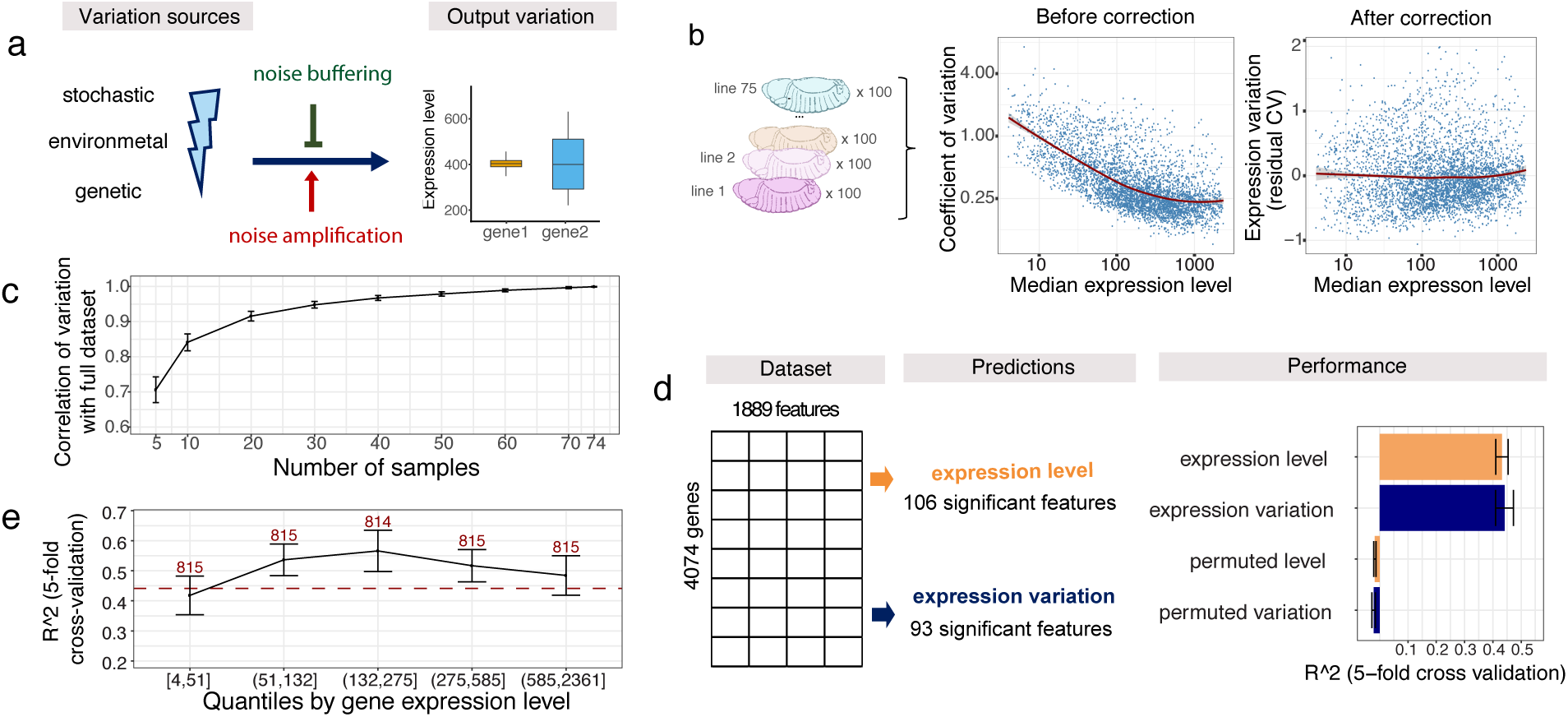
Genomic features can predict expression variation independent of expression levels. **(A)** Differences of gene regulatory mechanisms related to noise amplification and noise buffering would result in different observed expression variation given the same variation sources (left). **(B)** Dependence between coefficient of variation (CV) and median expression level of 4074 genes across 75 samples (left). Residuals from LOESS regression of CV on the median were used as the measure of variation throughout the analysis (right). Median expression level and coefficient of variation plotted on log2-scale, red line represents LOESS regression fit. **(C)** Correlation of expression variation calculated from subsets of samples versus the full data set. Error bars = standard deviation across 100 independent selections of samples. **(D)** Schematic overview of the random forest models and feature selection with Boruta algorithm (left). Performance shown as R^2 from 5-fold cross-validation and compared to randomly permuted data (right). Whiskers = standard deviation across the 5-fold cross validation. **(E)** Performance (R^2, 5-fold cross validation) for genes grouped by expression levels (quantiles). Whiskers represent standard deviation from 5-fold cross validation, number of genes per quantile indicated (x-axis). Red dotted line indicates performance of full model.

Over the last decade, studies on single-celled organisms or cell lines have linked multiple regulatory mechanisms to gene expression variation, including the presence of a TATA-box at the gene’s promoter (Ravarani et al. 2015; Blake et al. 2006), CpG islands (Morgan and Marioni 2018), bi-valent chromatin marks (Faure, Schmiedel, and Lehner 2017), polymerase pausing (Boettiger and Levine 2009) or miRNA binding (Schmiedel et al. 2015). However, it remains unclear what mechanisms regulate expression variation in multicellular, developing organisms in a gene and tissue-specific manner.

To address this, we devised a machine learning approach and performed a systematic analysis of factors underlying variation of gene expression in *Drosophila melanogaster* to uncover the regulatory mechanisms involved. To measure expression variation, we used gene expression data generated from a pool of embryos (∼100) sampled from 75 different isogenic lines during embryogenesis (Cannavò et al. 2016). This experimental design cancels out most stochastic noise (since it’s bulk sequencing), tissue-specific expression pattern (since it’s whole embryo) and slight differences in developmental progression (since it’s 100 embryos per line). To dissect the regulatory mechanisms that modulate expression variation (Fig 1a), we collated over a thousand gene-specific and genomically encoded features and applied a random forest model to identify the properties that best explain expression variation across individuals. As a comparison, we also predict median expression level across lines using the same features.

Our results show that, overall, increasing regulatory complexity translates into more robust gene expression. We identified two independent mechanisms associated with low expression variation across individual: Low variable genes either have (i) a broad transcription initiation region (broad promoters) with high transcription factor (TF) occupancy, or (ii) narrow initiation regions (narrow promoters) with Polymerase II (PolII) pausing and high regulatory complexity outside the promoter region. In contrast, genes with high variability generally have narrow promoters, and little other regulatory features, suggesting that it may rather be a lack of ‘stabilizing’ mechanisms that facilitates their noisy expression. Applying the same framework to human data derived from tissues across individuals (GTEx Consortium 2013) identified similar promoter-associated features to be predictive of expression variation, thus validating our findings in an independent organism. Remarkably, these same features are also predictive of differentially expressed genes when tested on independent datasets from adult *Drosophila* subjected to different stress conditions, or in a collection of differential expression data for human. These findings suggest that the differential expression response may be partially explained by genetically encoded gene-specific features that are unrelated to the treatment applied.

Taken together, our results suggest that gene expression variation across genetically diverse multicellular organisms is strongly linked to how the gene is regulated and likely reflects evolutionary constraints on expression precision.

## Results

### Measuring gene expression variation across individuals

To understand the mechanisms by which gene expression variation is controlled during embryonic development, we obtained RNA-seq data from 75 isogenic lines of *Drosophila melanogaster* embryos at three different developmental stages (2-4, 6-8, and 10-12 hours post fertilization) from (Cannavò et al. 2016). To reduce potential confounding effects of maternally deposited RNA, we focused on the late embryonic time-point (10-12 hours after fertilization), and removed genes whose expression decreased between 2-4 h and 10-12h, resulting in embryonic expression data for 4074 genes (Methods, Supplementary Fig 1). For each gene, we calculated its median expression level and the coefficient of variation (CV) from the normalized read counts across individuals (Methods). As variation is highly correlated with the levels of gene expression (Anders and Huber 2010; Ran and Daye 2017; Eling et al. 2018) we used the residuals from a locally weighed regression (LOESS) of the CV on median expression to obtain a measure of expression variation that is relative to the expected variation at a given expression level (Fig. 1b).

We confirmed that this measure of variation is highly correlated with alternative metrics, such as variance stabilized standard deviation or residual median absolute deviation (Supplementary Fig. 2a-b) and robust with respect to the number and identity of samples used (Fig. 1c). Moreover, using the full dataset from Cannavò (Cannavò et al. 2016), expression variation values were highly correlated across time, especially for consecutive time-points, further confirming the approach (Methods, Supplementary Fig.1d). Finally, we observed a strong correlation in expression variation between pairs of genes in close proximity (Supplementary Fig. 1e), as previously observed for neighbouring genes in yeast (Becskei, Kaufmann, and van Oudenaarden 2005; Batada and Hurst 2007).

As these 75 samples came from strains with different genotypes, we first calculated the proportion of expression variance that is explained by genetics in *cis* (taking variants within 50 kb of each gene into account) using variance decomposition (Methods). On average, 6% (median across all genes) of the total gene expression variation was explained by cis genetics (Supplementary Fig.1f), indicating that more complex genetic effects and other properties must account for the majority of expression variation. We reasoned that differences in the extent of expression variation among genes should reflect inherent differences in their regulation, including their regulatory complexity and mechanisms of noise buffering or amplification. Therefore, in the remainder of this study we investigate the regulatory differences between genes with high versus low expression variation.

### Genomic features predict expression variation independent of expression levels

To understand the drivers of expression variation, we collected 1,888 gene-specific features (Table 1, Supplementary tables 1-3) and used random forest regression to identify those that are associated with either expression variation or expression level (Fig 1d). This allowed us to distinguish between features that are predictive of one or both properties. The features can be broadly divided into seven categories: transcription start site (TSS; e.g. core promoter motifs, chromatin accessibility, TF binding), gene body features (e.g. gene length, number of exons), 3’untranslated regions (UTR; e.g. length, miRNA motifs), distal regulatory elements (e.g. TSS-distal chromatin accessibility, TF occupancy), gene type (e.g. housekeeping genes, TFs), gene context (e.g. gene density, distance to the borders of topologically associated domains (TADs)), and genetics (e.g. the presence of eQTL and a *cis* genetic component; full description in Methods and Table 1).

**Table 1.**
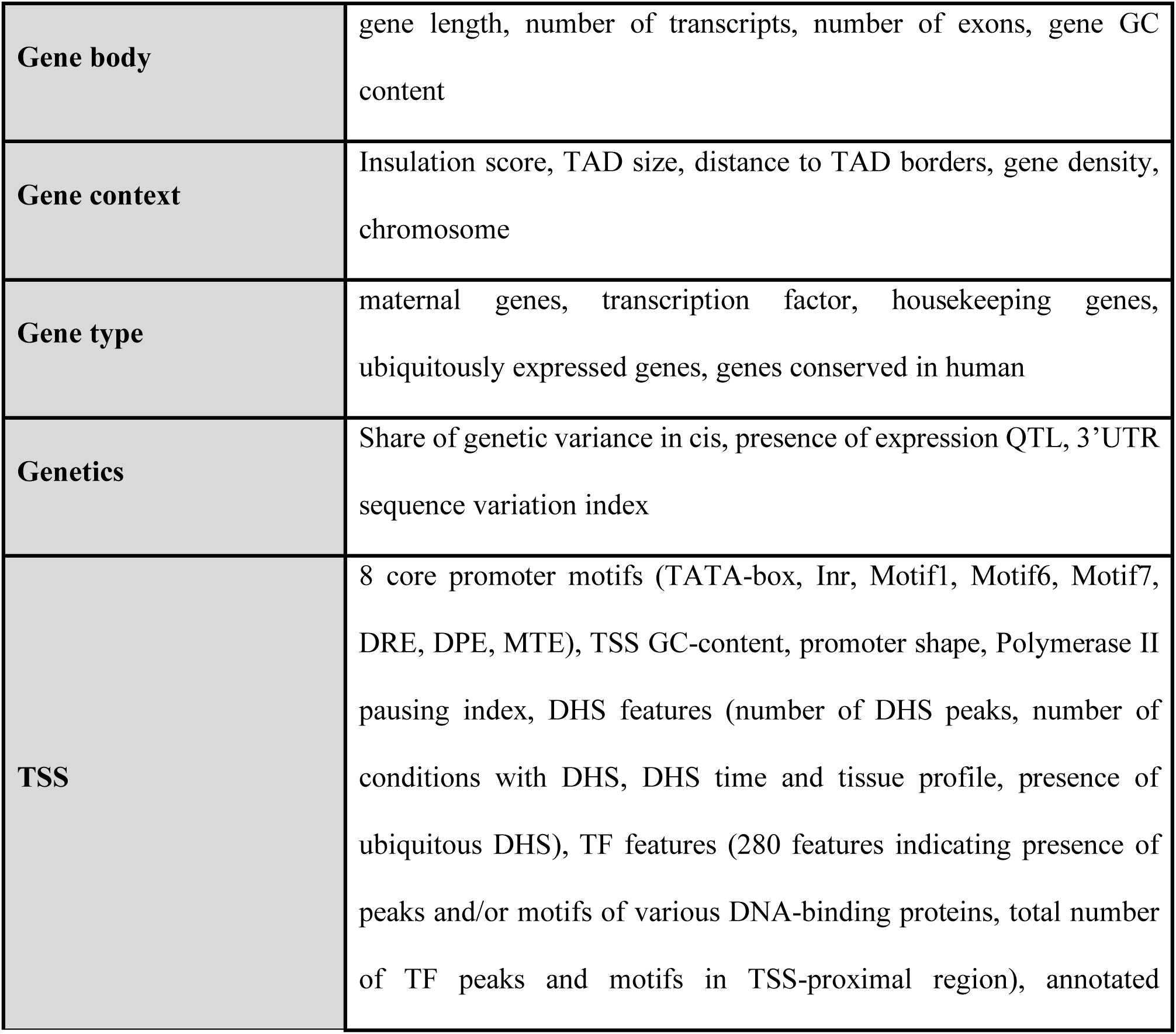
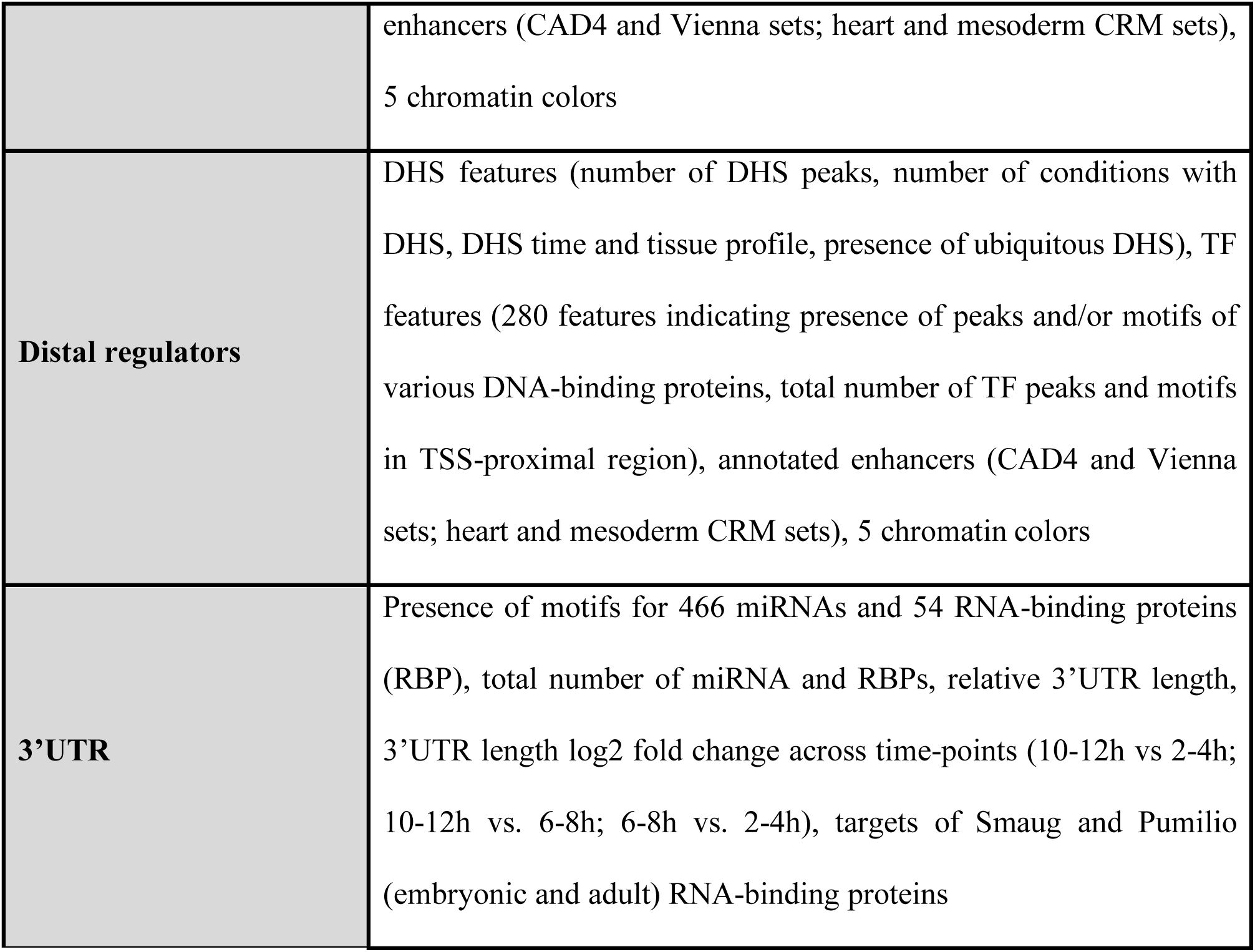
Summary of features in Drosophila table by class. Description of features and sources are provided in methods.

|  |  |
| --- | --- |
| **Gene body** | gene length, number of transcripts, number of exons, gene GC content |
| **Gene context** | Insulation score, TAD size, distance to TAD borders, gene density, chromosome |
| **Gene type** | maternal genes, transcription factor, housekeeping genes, ubiquitously expressed genes, genes conserved in human |
| **Genetics** | Share of genetic variance in cis, presence of expression QTL, 3'UTR sequence variation index |
| **TSS** | 8 core promoter motifs (TATA-box, Inr, Motif1, Motif6, Motif7, DRE, DPE, MTE), TSS GC-content, promoter shape, Polymerase II pausing index, DHS features (number of DHS peaks, number of conditions with DHS, DHS time and tissue profile, presence of ubiquitous DHS), TF features (280 features indicating presence of peaks and/or motifs of various DNA-binding proteins, total number of TF peaks and motifs in TSS-proximal region), annotated |
|  | enhancers (CAD4 and Vienna sets; heart and mesoderm CRM sets),<br>5 chromatin colors |
| **Distal regulators** | DHS features (number of DHS peaks, number of conditions with DHS, DHS time and tissue profile, presence of ubiquitous DHS), TF features (280 features indicating presence of peaks and/or motifs of various DNA-binding proteins, total number of TF peaks and motifs in TSS-proximal region), annotated enhancers (CAD4 and Vienna sets; heart and mesoderm CRM sets), 5 chromatin colors |
| **3'UTR** | Presence of motifs for 466 miRNAs and 54 RNA-binding proteins (RBP), total number of miRNA and RBPs, relative 3'UTR length, 3'UTR length log2 fold change across time-points (10-12h vs 2-4h; 10-12h vs. 6-8h; 6-8h vs. 2-4h), targets of Smaug and Pumilio (embryonic and adult) RNA-binding proteins |

To restrict our analysis to the important features, we applied the random forest-based Boruta algorithm, which iteratively selects all features that predict better than their permuted version (Kursa and Rudnicki 2010). This resulted in 93 and 106 predictive features for expression variation and level, respectively (Fig. 1d). Using these feature sets, our models predicted expression variation and level with an R^2^ of 0.45 and 0.43 (5-fold cross validation), respectively, while permuting the labels resulted an R^2^ of zero (Fig. 1d).

To ensure the robustness of our predictions we have performed a number of analyses: first, we verified that the predictions for variation are independent of the level of gene expression by showing that the models performed equally well on genes grouped into quartiles based on their expression levels (Fig. 1e). Second, we ensured that the predictions are robust to the choice of measure used for expression variation (Supplementary Fig. 2c). Third, we tested whether dynamic gene expression changes during developmental stages can contribute to the variation predictions. We reran the random forest models, predicting expression variation for genes grouped based on their absolute expression change between 6-8 and 10-12 hours after fertilization. For genes with minor expression change between the two time-points (below median of 0.8), the performance was comparable to the full model, while for the genes with a stronger expression change (above 0.8) the R^2^ dropped to about 0.3 (Supplementary Fig. 2d). This indicates that some portion of expression variation comes from dynamic changes in gene expression during embryogenesis, which is not captured by our features (and thus reduces the performance of our model for this set of genes). However, since the performance is the best for genes that vary little between stages, it indicates that variance explained by our model is overall not majorly confounded by expression dynamics. Finally, the model performance does not decrease when training and test sets come from different chromosomes (Supplementary Fig. 2e), demonstrating that the results are not confounded by shared regulatory features between neighboring genes.

Taken together, these results establish that gene expression variation - as well as gene expression levels - can be predicted based on genomically encoded features, when measured across a population of genetically diverse individuals during embryogenesis. The predictions are independent of the gene’s expression level and are robust to the metric used for measuring variation. These models can therefore be used as the basis for addressing questions about buffering mechanisms that regulate gene expression variation during embryogenesis.

### Promoter architecture is the most important predictor of expression variation

Next, we used this predictive framework to investigate the genomic features that best explain expression variation and expression level. We retrieved the features’ ‘importance score’ from the Boruta algorithm and determined the correlation of each feature with both expression properties (Supplementary Table 4). Although most features are to some extent predictive of both expression level and variation, their relative importance differed substantially (Fig. 2a). Being a housekeeping gene, for example, was strongly predictive of high expression level while being less important for expression variation. Conversely, the presence of a core promoter TATA-box motif is strongly predictive of high expression variation only (Fig. 2a, see Suppl. Table 4 for full list). We note that most features are either associated with higher variation and lower expression or vice versa, suggesting that expression level and variation are not completely independent, as was previously observed (Faure, Schmiedel, and Lehner 2017), even though they are globally uncorrelated (Fig 1b). However, we found that when we split genes into the categories of the top features (e.g. housekeeping vs non-housekeeping) the differences in expression variation are pronounced at all expression levels (Fig 2b-e): For example, housekeeping genes (the strongest predictor for expression level) are less variable than non-housekeeping genes at any level of expression (Fig 2b). The same holds true for the feature ‘promoter shape index’, which is the strongest predictor for variation (Fig 2c), as well as other features such as ‘#conditions with DHS’ (DNase hypersensitive sites) (Fig 2d) and ‘presence of a TATA box’ (Fig 2e). This demonstrates that the features explain expression variation independent of expression level.

**Figure 2.**
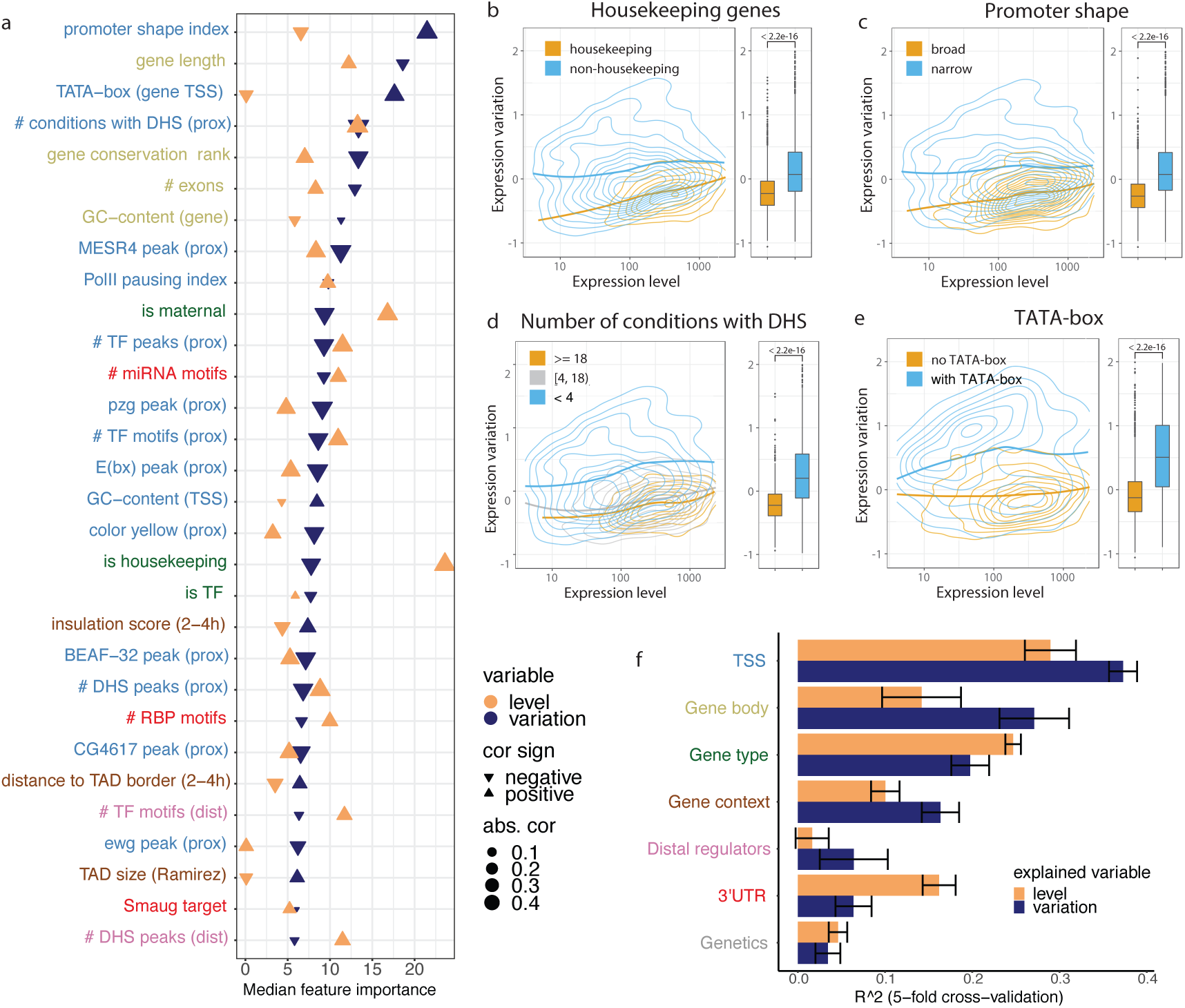
Promoter architecture is the most important predictor of expression variation. **(A)** Top-30 important features for predicting expression variation using Boruta feature selection. Features are ordered by their importance for expression variation (blue) and show the corresponding importance for level (orange). The absolute value and sign of correlation coefficient is indicated by the triangle size and orientation, respectively. For binary features, phi coefficient of correlation was used, otherwise Spearman coefficient of correlation. Label colors correspond to feature groups in (F). **(B-E)** Relationship between expression level and expression variation shown as 2D kernel density contours (left) and boxplots (right) for housekeeping genes **(B)**, genes separated by promoter shape **(C)**, number of embryonic conditions with a DHS **(D)**, and presence of TATA-box at TSS **(E)**. LOESS regression lines indicated for each gene group, P-values from Wilcoxon test. **(F)** Performance of random forest predictions (mean R^2 from 5-fold cross-validation) for expression level (orange) and variation (blue) trained on individual feature groups. Whiskers = standard deviation, color code of y-axis labels matches Fig 2A.

Promoter-associated features (*TSS-proximal*) are among the strongest predictors in terms of explanatory power for expression variation, and include promoter shape, core promoter motifs and GC-content, Pol II pausing, chromatin accessibility, and TF occupancy at TSS (Fig. 2a). Consequently, a model based only on TSS-proximal features can predict expression variation fairly well with R^2^=0.37, while performing less well for predicting expression level (R^2^= 0.29) (Fig. 2f). Although lower than the model using all features (R^2^ of variation/level 0.45/0.43), this is markedly higher than a model on any other feature type alone. The next most predictive class of features for variation are *gene body* (R^2^ 0.27/0.14) and *gene type* (0.20/0.25) (although more predictive of expression level), followed by *gene context* (0.16/0.10). *3’UTR* features, which rank third among the most predictive features of expression levels, show little predictive value for variation (0.06/0.16), and distal features overall showed a rather weak predictive value for both variation and level (0.06/0.01). Finally, *Genetics* was the least predictive for both variation and level among the seven feature groups (0.03/0.05), in keeping with the variance decomposition analysis above.

In summary, our results demonstrate that multiple regulatory features can independently predict gene expression variation or gene expression levels. Interestingly, promoter features, rather than upstream regulatory complexity (such as distal DHS sites), are the most predictive of expression variation. Given that housekeeping genes and TFs tend to have different promoter types (Arnold et al. 2016; Haberle and Stark 2018; Lenhard, Sandelin, and Carninci 2012), this suggests that specific biological functions may have distinct mechanisms to reduce variation and provide robustness to their expression as evidenced by models based solely on a gene’s functional annotation (*Gene type* in Fig. 2f).

### Expression variation in broad versus narrow promoter genes reflects trade-off between expression robustness and plasticity

The most prominent predictive feature for expression variation is promoter shape index (Fig 2a), which classifies promoters based on the broadness of their transcriptional initiation region (Schor et al. 2017; Rach et al. 2009; Forrest et al. 2014; Lenhard, Sandelin, and Carninci 2012). Genes with narrow promoters generally have higher variation compared to genes with broad promoters (Fig. 2c), and, interestingly, also comprise a wider range of variation (Fig 3a). Moreover, expression variation of narrow promoter genes is better explained by genomically encoded features compared to broad promoter genes (R^2^= 0.37 vs 0.14), and this difference in performance becomes more pronounced with more stringently defined narrow and broad promoter genes (Fig. 3b).

**Figure 3.**
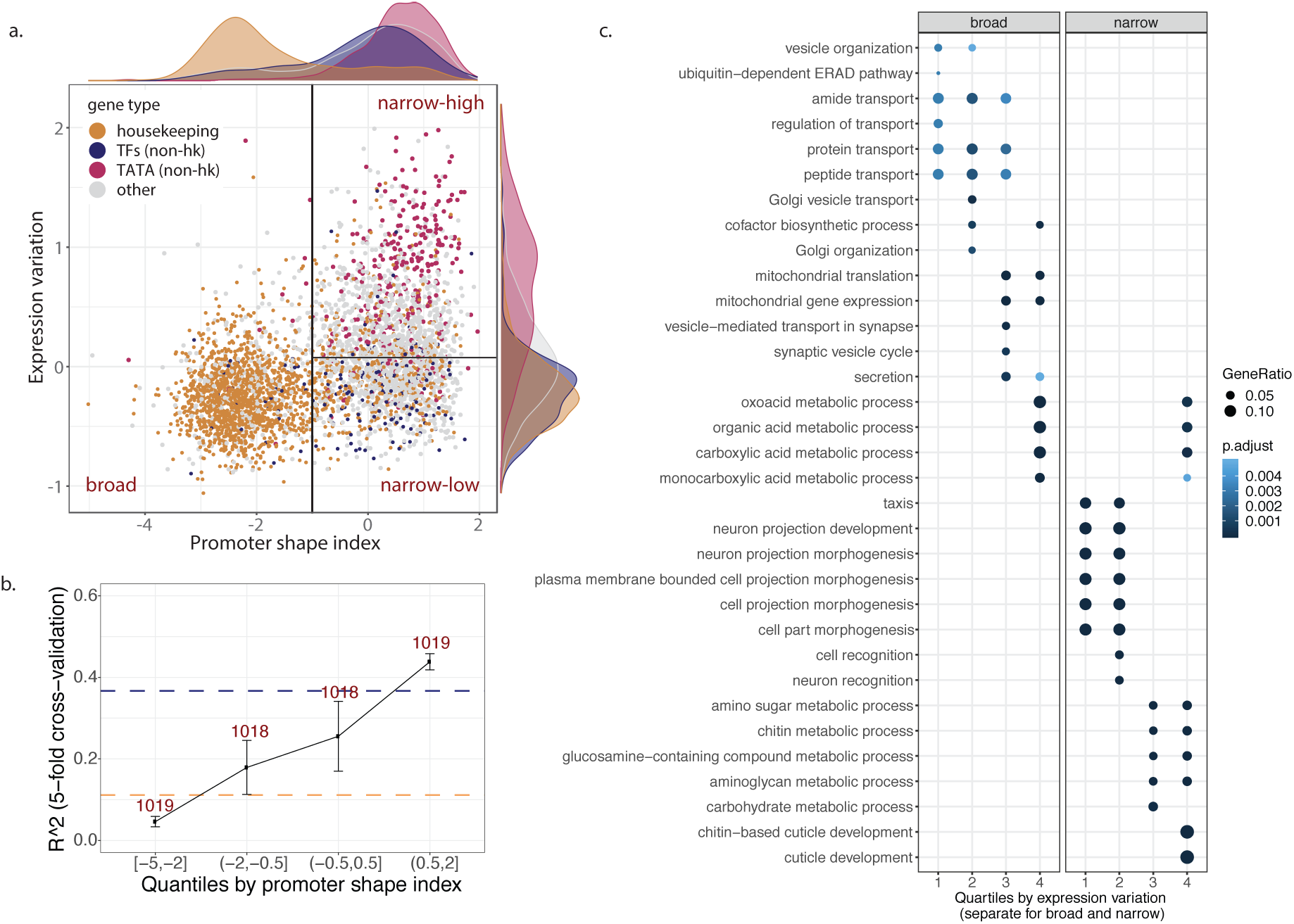
Expression variation in broad versus narrow promoter genes reflects trade-offs between expression robustness and regulatory plasticity. **(A)** Genes separate into three groups based on their promoter shape index (x-axis) and expression variation (y-axis). Each dot represents a gene; colors indicate gene annotations: housekeeping (orange), non-housekeeping TFs (blue), non-housekeeping with a TATA-box (red), other (grey). Distributions of promoter shape index and expression variation across gene groups are shown as density plots. Broad and narrow promoter genes are separated based on shape index threshold of −1 (vertical black line) as in (Schor et al. 2017). Narrow-low and narrow-high groups are separated based on the median expression variation of narrow promoter genes (horizontal black line). **(B)** Performance to predict expression variation for genes split by quartiles of promoter shape index. Horizontal lines show performance (mean R^2 from 5-fold cross-validation) on broad (orange) and narrow (blue) promoter genes separately. Whiskers = standard deviation (from 5-fold cross validation), number of genes per categories indicated (x-axis). **(C)** GO term enrichment (Biological Process) of genes stratified by promoter shape and expression variation. Top GO terms are shown (full list in Supplementary Table 6. Quartiles of expression variation (1-lowest, 4 – highest) were calculated for broad and narrow promoter genes separately. Quantile intervals for broad and narrow promoter genes provided in methods.

Interestingly, when we group genes from the two promoter classes into quartiles based on their variation we find very specific functions enriched among them: the broad class is strongly enriched for housekeeping genes (Fishers’s test odds ratio, OR=15.0, p-value<1e-16, Supplementary table 5) and GO terms related to basic cellular processes (cellular transport, secretion, and DNA/RNA biogenesis) with the exception of the top 25% most variable genes within the group being also enriched in metabolic processes (Fig. 3c, Supplementary Fig. 3a, Supplementary Table 6). In contrast, narrow promoters genes fall into two functional categories depending on their expression variation: the bottom 50% were enriched in TFs (OR=3.0, p-value<1e-16) and GO terms related to development, signaling and regulation of transcription, while the top 50% are enriched for TATA-box genes (OR=7.9, p-value<1e-16) and GO terms related to metabolism, stress response, and cuticle development (Fig. 3c, Supplementary Fig. 3a). We therefore grouped genes along the dimensions of promoter shape and expression variation into three classes (Fig. 3a): genes with broad promoters and low levels of variation in expression (broad), genes with narrow promoters and low expression variation (narrow-low) and genes with narrow promoters and high expression variation (narrow-high).

Next we looked at regulatory plasticity of these classes of genes defined here as the variation in accessibility of TSS-proximal DHSs across time and tissues, see [Reddington et al, submitted]. We observed that narrow promoter genes had high regulatory plasticity regardless of their expression variation (Supplementary Fig. 3b). In particular, narrow-low genes are robustly expressed across individuals at the given developmental stage, while having condition-specific regulation. In contrast, broad promoter genes are characterized by both low expression variation and low plasticity, which agrees with their housekeeping functions.

Enrichment of low-variable genes in either housekeeping (*broad*) or developmental (*narrow-low*) functions suggests selection pressure may act on those genes to reduce expression noise in genes essential for viability and development. One proxy for evolutionary constraints is sequence conservation across long evolutionary distances. In keeping with this, sequence conservation between *Drosophila* and human was among the top five most predictive features of low expression variation with conserved genes being significantly less variable (Fig 2a, Wilcoxon test p-value <2e-16). Promoter shape is also correlated with gene conservation: conserved genes are highly enriched for broad promoters (80% in broad vs. 41% in narrow) and more enriched in the narrow-low compared to narrow-high class (54% vs 28%). Within each class, conserved genes are less variable (Supplementary Fig.3c), hence sequence conservation provides additional information about variation constraints across genes.

Overall, these results suggest that expression variation is an orthogonal component to the regulatory plasticity, which has previously been defined along the narrow-broad promoter spectrum (Rach et al. 2009; Lenhard, Sandelin, and Carninci 2012). Promoter shape likely reflects differences in regulatory plasticity (constitutive vs. condition-specific genes), while expression variation may reflect evolutionary constraints on expression robustness with essential and highly conserved genes being less variable. These findings indicate a partial uncoupling between expression variation across multicellular individuals in a controlled environment and variation across tissues/development, analogous to the uncoupling between plasticity and noise observed in yeast (Lehner 2010), and suggest different mechanisms to control expression robustness for genes with ubiquitous versus condition-specific expression.

### Two classes of genes with low variation have distinct regulatory mechanisms

The results above indicate that the partial uncoupling of expression variation and expression plasticity could be achieved by distinct mechanisms of ensuring expression robustness between different promoter architectures (broad/narrow). To explore this, we examined the most predictive features in relation to the different promoter types. Among the most significant promoter features is “#conditions with DHS” (Fig 2a), which is derived from a comprehensive tissue and embryonic stage specific atlas of open chromatin regions (DHS data for 19 conditions) during a time-course of *Drosophila* embryogenesis (Reddington et al, submitted). The median number of developmental conditions in which a gene had at least one DHS site was 18, 8, and 1 for broad, narrow-low, and narrow-high genes respectively (Fig 4a), thus highlighting again that the narrow-low and broad classes differ in their developmental plasticity (Fig. 4a). A similar trend was observed for related features, such as using a compendium of TF occupancy data during embryogenesis (Fig 4b), TF peaks with motifs, or motifs alone (Supplementary Fig.4a-b). To understand how these promoter-type specific DHS patterns are set-up we next examined the 24 TFs that were predictive of expression variation in the full model (Supplementary Table 4, *‘med_imp_var’* column). Broad promoter genes were generally strongly enriched for ubiquitously expressed TFs, insulator proteins and chromatin remodelers (e.g. BEAF-32, MESR4, E(bx); Fig 4c, Supplementary table 5; Fishers exact test). The narrow-low class was enriched for the Polycomb-associated developmental factors Trl and Jarid2, while the narrow-high were not strongly enriched for any TF (Fig. 4c). Interestingly, some of the TFs enriched in broad vs narrow promoters, are still predictive of expression variation in the narrow-promoter only model (e.g. MESR4, E(bx), and YL-1, Supplementary Fig.4c), while the presence of ‘narrow’ TFs, despite being associated with low variation in narrow promoters, had the opposite effect in the broad class (Fig. 4c bottom right).

**Figure 4.**
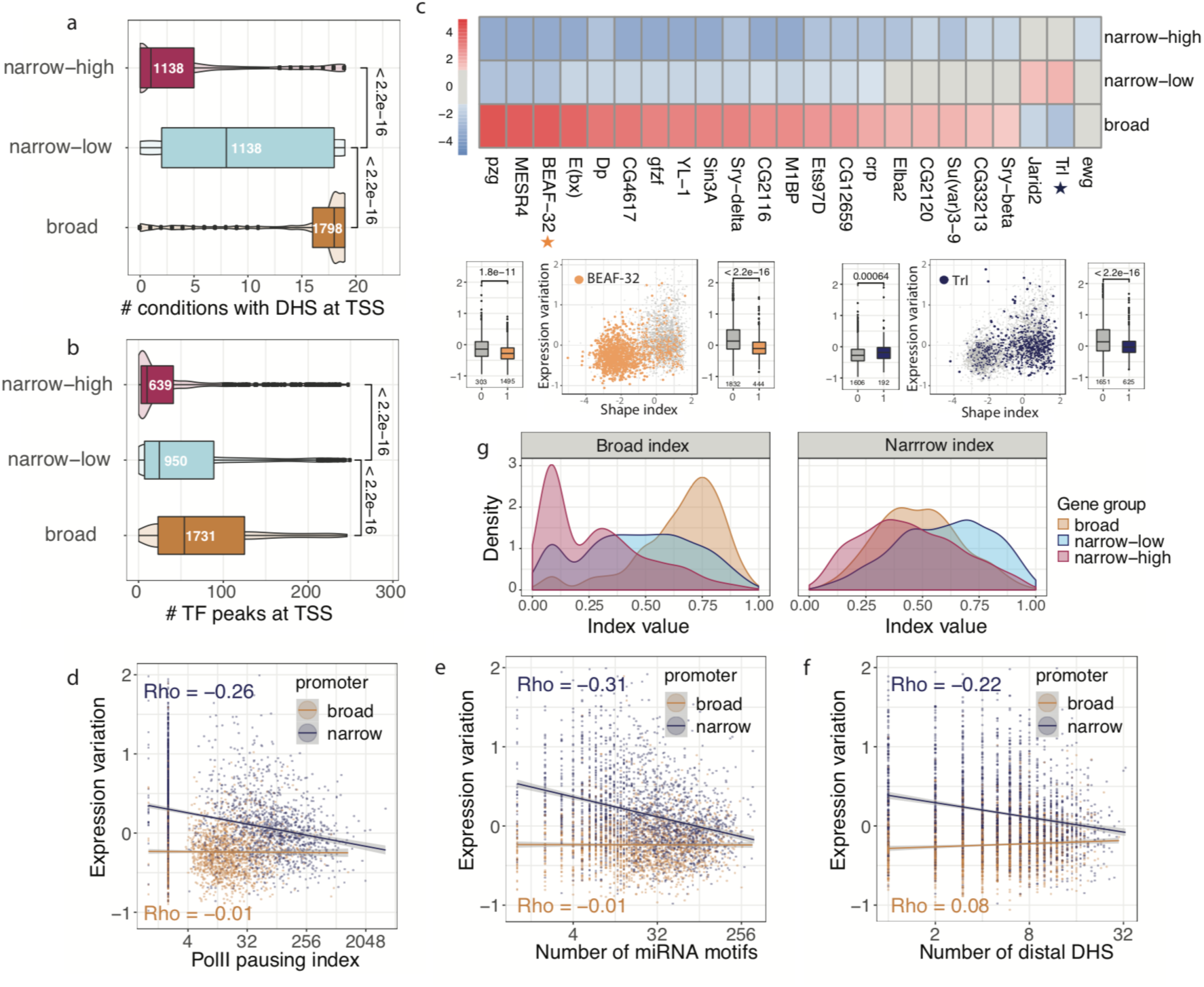
Different regulatory mechanisms lead to expression robustness in genes with broad and narrow promoters. **(A,B)** Chromatin accessibility (number of conditions with DHS) **(A)**, or number of different TF peaks **(B)** overlapping TSS-proximal DHS for genes stratified into broad, narrow-low and narrow-high (defined in Fig 3A). P-values from Wilcoxon test. **(C)** Top: enrichment (odds ratio from Fisher’s test) of ChIP peaks for 24 TFs in TSS-proximal DHSs of broad, narrow-low and narrow-high genes. Only TFs with predictive importance for expression variation (based on Boruta) were included. For each TF, Fisher’s test was performed separately for each category vs all other. Color = log2 odds ratio from Fisher’s exact test (two-sided), grey = non-significant comparisons (adjusted p-value cutoff of 0.01, Benjamini-Hochberg correction on all 24×3 comparisons). Lower panels: Presence of BEAF-32 (left) and Trl (right) ChIP-seq peaks in TSS-proximal DHS, plotted coordinates of promoter shape index and expression variation (same as Fig. 3a). Each dot represents a gene (grey if TF peak is absent, blue for Trl, orange for BEAF-32). **(D-F)** Relationship between polymerase pausing index **(D)**, number of miRNA motifs in 3’UTR of a gene **(E)** and number of TSS-distal DHS peaks **(F)** and expression variation for broad (orange) and narrow (blue) promoter genes. Each dot represents a gene, lines linear regression fits, rho=Spearman correlation coefficient. **(G)** Gene scores by two indices constructed as the normalized rank average of: number of embryonic conditions with DHS, number of TF peaks, number of TF motifs (Broad regulatory index;; left), and number of TSS-distal DHS, number of miRNA motifs, Pol II pausing index (Narrow regulatory index; right). Colors correspond to broad (orange), narrow-low (blue) and narrow-high (red)) gene groups. P-values < 1e-09 for all pairwise comparisons of the distributions.

The next most predictive feature in our model is “PolII pausing index” (Fig 2a), defined as the density of polymerases in the promoter region divided by the gene body length (Saunders et al. 2013)(Fig 2a). Narrow-low genes have the highest pausing index (40) followed by broad and narrow-high genes (10 and 7, respectively; Supplementary Fig.4d). Consequently, Pol II pausing is strongly negatively correlated with expression variation in narrow promoters (Spearman correlation Rho=-0.28, p-value<1e-16), yet showed no significant relationship in broad (Fig. 4d), again highlighting different mechanisms to confer robust expression. This may be partially explained by Trl, which can modulate the level of Pol II pausing (Tsai et al. 2016).

Among the most significant non-promoter features, our model identified distal regulatory complexity (“#TF motifs (dist)” and “#DHS peaks (dist)” in Fig 2a) and post-transcriptional events (“#miRNA motifs” and “#RBP motifs” in Fig 2a) as predictive of expression variation. As for the distal regulatory complexity, narrow-low had the highest number (median of 6) of distal regulatory elements, defined as DHS within 10kb of the TSS, followed by broad (4) and narrow-high (4) genes (Supplementary Fig.4g). Consequently, the number of distal DHS is negatively correlated with expression variation in narrow promoters (Rho=-0.22, p-value<1e-16) while being uncoupled from variation for broad (Fig. 4f). Similarly, narrow-low genes have a higher number of miRNA motifs in their 3’UTRs (median of 35) compared to broad (20) and narrow-high (14) genes (Supplementary Fig.4e), which again was negatively correlated with variation in narrow promoter genes only (Rho=-0.31, p-value<1e-16) (Fig. 4e). Similar results were obtained for the number of RNA-binding protein (RBP) motifs, which have an effect for narrow, but not for broad, genes (Supplementary fig. 4f).

In summary, these findings provide strong evidence that robustness in gene expression across individuals is conveyed by different mechanisms depending on the gene’s promoter type: in broad promoter genes, robust expression is likely a result of a plethora of broadly expressed TFs that bind to the core promoter and keep the chromatin constitutively accessible, compatible with their house-keeping roles. Narrow promoter genes, in contrast, seem to be regulated by a smaller number of (narrow-specific) TFs and their robustness is conveyed through mechanisms that involve Pol II pausing, distal regulatory elements, and posttranscriptional regulation. This suggests that broad and narrow promoter types have distinct mechanisms to regulate expression variation that are not necessarily transferable. This is possibly related to the relatively higher regulatory plasticity required of the narrow-low genes.

Partial aspects of these findings have been reported previously. E.g. In a study of 14 developmental control genes, Pol II pausing at promoters was linked to more synchronous gene activation, thereby reducing cell-to-cell variability in the activation of gene expression (Boettiger and Levine 2009). Also, miRNAs have been proposed to buffer expression noise (Schmiedel et al. 2018, 2015). Our data puts these previous findings in a more global context as part of a distinct mechanism for a particular promoter type.

We summarized these mechanisms as two indices based on the ranked averages of the corresponding features: *broad regulatory index* (number of TF peaks, motifs and conditions with DHS, at the TSS) and *narrow regulatory index* (Pol II pausing index, number of distal DHS and miRNA motifs), respectively (Fig 4g), which nicely separate the three gene groups. Interestingly, we found no evidence for a specific noise-amplifying factor, except for the TATA-box. Yet, even for TATA-box genes, since they are depleted of all the aforementioned robustness features (Supplementary Fig. 4h), the observed high variation may result from a lack of robustness-conveying factors.

### Expression variation can predict signatures of differential expression upon stress

So far, we showed that distinct mechanisms regulating expression variation are directly encoded in the genome. In the following, we want to assess the impact of these findings for interpreting gene expression studies in general.

We postulate that the expression variation of a gene across individuals can be interpreted as its ability to be modulated by any random perturbation. If this is true, we expect expression variation to be predictive of a gene’s response to changes in the environment. To test this, we used an independent gene expression dataset from adult flies that were subjected to different stress conditions related to temperature, starvation, radiation, and fungi infection (Moskalev et al. 2015). In agreement with our postulation we find that genes differentially expressed upon stress have high expression variation in our embryonic dataset (Fig. 5a, Wilcoxon test p-value<1e-16). Remarkably, this held true for every individual stress condition (Supplementary Fig. 5a).

**Figure 5.**
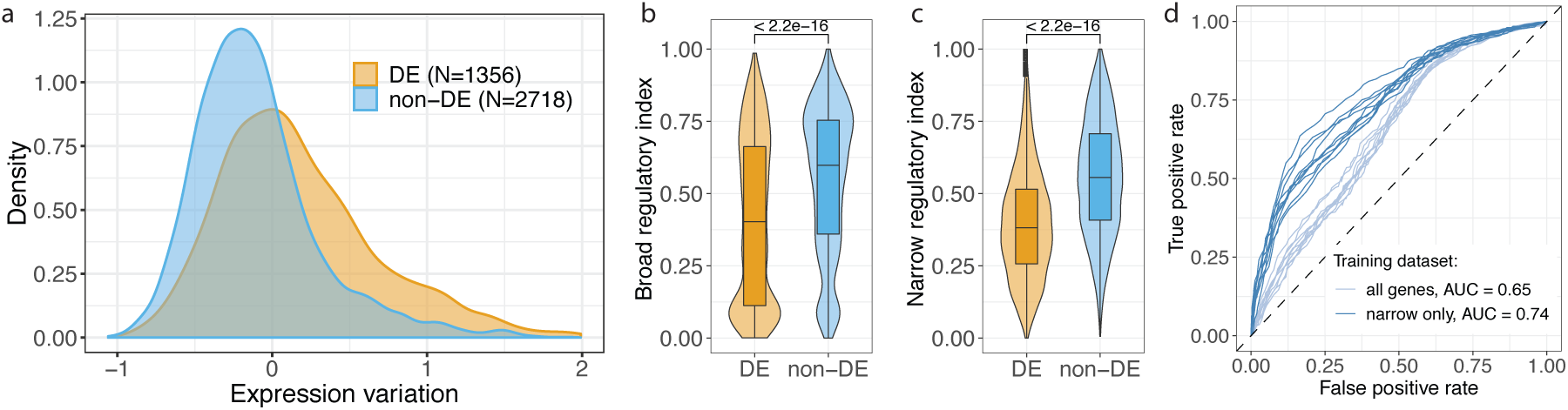
Expression variation can predict signatures of differentially expression upon stress. **(A)** Expression variation of genes differentially expressed (DE) upon any stress conditions from (Moskalev et al. 2015) compared to non-differentially expressed genes (non-DE). **(B-C)** Differences in scores by the regulatory complexity indices (from Fig. 4g) between DE and non-DE genes (from Fig. 6a): broad regulatory index **(B)**, narrow regulatory index **(C)**, P-values from Wilcoxon rank test. **(D)** ROC-cures for predicting DE with random forest models trained on expression variation (top-30% variable vs. bottom-30% variable) in all genes (light blue) or narrow promoter genes (dark blue). Models were trained and tested on non-overlapping subsets of genes in 10 random sampling rounds (all plotted). Median AUC values from 10 sampling rounds.

Differentially expressed genes are enriched for narrow-high promoter genes (Fishers’s test odds ratio=2.97, p <1e-16). Consequently, they are associated with lower chromatin accessibility (p <1e-16, Supplementary Fig. 5b), a lower number of TFs (p=1.4e-10) and less motifs (p=3.9e-8) at their TSS, as well as other features important for distinguishing between narrow-high and -low genes (Supplementary Fig. 5c-e). Overall, differentially expressed genes showed lower regulatory complexity as reflected in our broad and narrow variability indices (Fig 5b-c).

To assess this association more systematically, we next tested whether the model for predicting expression variation can also identify differentially expressed genes. We trained a random forest model using our embryonic data to classify the top-30% versus bottom-30% variable genes and used it to predict differential expression in adults subjected to different stresses (Methods). The model predicted differential expression on the non-overlapping test set with an AUC of 0.65 and 0.74 when trained to predict embryonic variation for all genes, or for narrow promoter genes, respectively (Fig. 5d). This demonstrates that differential expression can be predicted based on a model trained for predicting expression variation. Since the model’s performance was better when trained only on variation in narrow promoters, it is likely that the narrow-specific regulatory mechanisms, such as micro RNA and enhancers, determine a gene’s responsiveness to stress. This is also reflected by the strong differences in narrow index between DE and non-DE genes (Fig 5c).

Overall, this suggests that the same buffering mechanisms confer expression robustness to different kinds of perturbations. Since the propensity to be differentially expressed is predictable based on genomically encoded features, this implies that results from differential expression studies should always be interpreted relative to a genes inherent tendency to respond to perturbation.

### Human promoter features predict both expression variation and differential expression

Given that gene expression variation across individuals can be predicted from genomic features in *Drosophila* we next asked whether this holds true in humans, and whether the predictive features are conserved. We used high quality RNA-seq datasets from the GTEx project comprising 43 tissues with data for at least 100 individuals (GTEx Consortium 2013). For each tissue, we measured expression variation across individuals using the coefficient of variation corrected for mean-variance dependence, applying a similar approach as for *Drosophila* (Methods). Since gene expression variation values were highly correlated across all tissues (Supplementary Fig 6), we also computed the mean of tissue-specific variations (mean variation) as potentially more robust metrics.

Since TSS-proximal features were the most predictive of expression variation in fly, we focused on promoter features to train the models (Methods). This included promoter shape, TF binding at the TSS, chromatin states, and several sequence features (TATA-box, GC-content, CpG islands). To predict the mean expression variation, promoter shape and chromatin state features were aggregated across multiple tissues. In addition, we collated three tissue-specific datasets for muscle, lung and ovary by matching RNA-seq, CAGE and chromHMM datasets (Methods). Based only on these features, random forest models were able to predict expression variation and level within each tissue to a similar extent as in *Drosophila* embryos (Fig. 2f) with R^2 ranging between 0.38-0.46 for expression variation and 0.19-0.24 for expression level (Fig. 6a). Aggregating expression variation across tissues yielded even higher performance with R^2 of 0.56 versus 0.31 for mean level across all expressing tissues. The overall performance was robust to changes in the numbers of samples including subsetting by age or sex (Supplementary fig. 8a).

**Figure 6.**
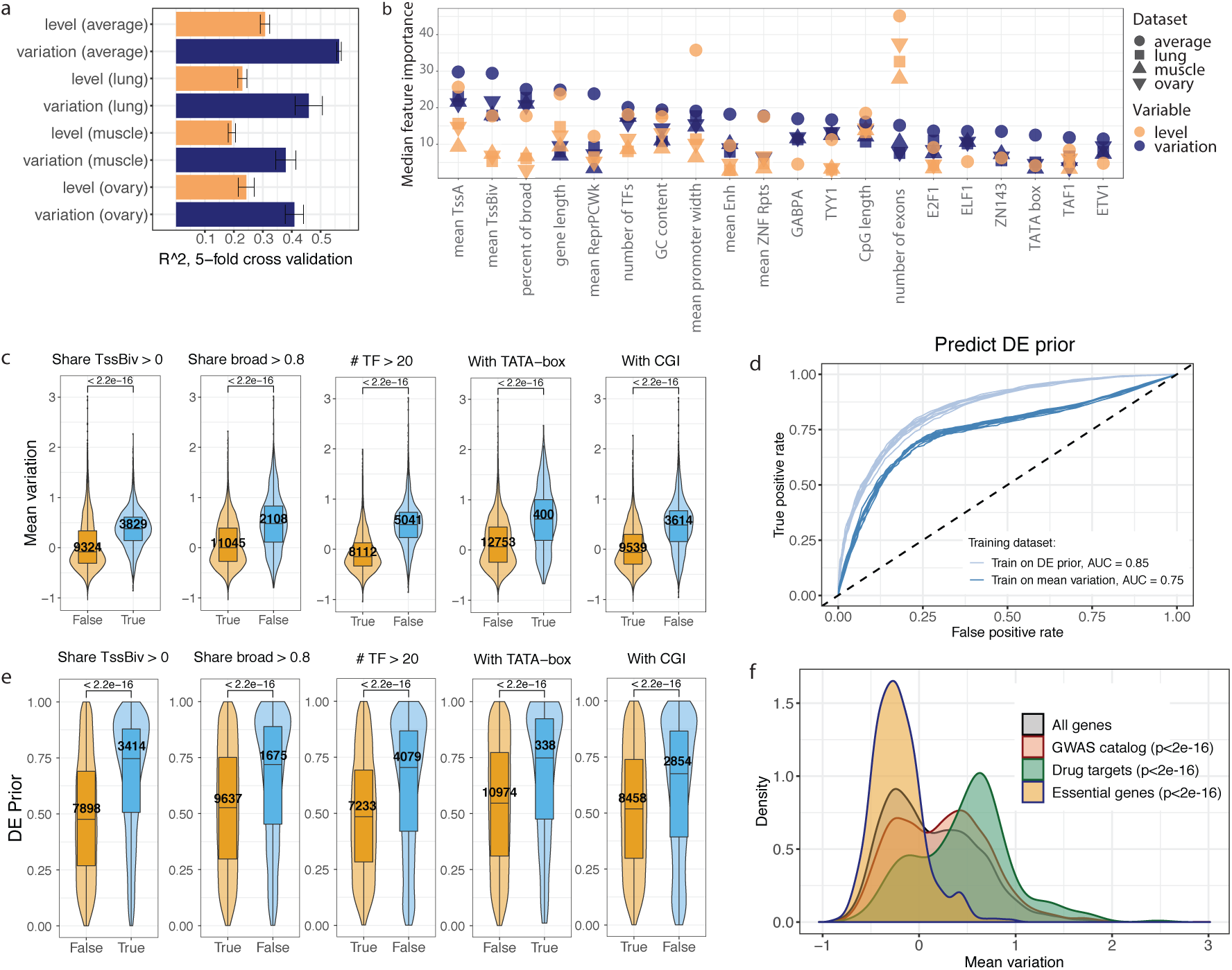
Features in human promoters predict both expression variation and differential expression. **(A)** Performance of random forest predictions (mean R^2 from 5-fold cross-validation, whiskers = standard deviation) for expression level (orange) and variation (blue) trained on expression variation in tissue-specific RNA-seq (lung, ovary, and muscle), as well as mean variation across 43 tissues (Methods). **(B)** Top-20 features for predicting expression variation using Boruta feature selection. Features ordered by their importance for expression variation (blue), showing the corresponding importance for level (orange). Shapes indicate four different datasets (three tissues and mean variation). **(C,E)** Differences in expression variation (C) and DE prior (E) for some of the top-predictive features from (B). P-values = Wilcoxon test, number of genes indicated. ‘Share TssBiv > 0’ indicates genes that have “TSS bivalent” chromatin state (chomHMM, Methods) in at least one tissue. ‘Share broad > 0.8’ indicates genes which have broad promoter in at least 80% of tissues where it is expressed (Methods). (D) ROC-curves for predicting DE prior (top-30% variable vs. bottom-30%) with random forest models trained on DE prior (light blue) and mean expression variation (dark blue). Models trained and tested on non-overlapping subsets of genes in 10 random sampling rounds (all plotted), with median AUC values indicated. **(F)** Mean expression variation of specific genes groups (GWAS hits, essential genes, drug targets) compared to the distribution of mean expression variation for all genes in the dataset.

The predictive features of expression variation in humans are highly overlapping with those for *Drosophila* (Fig 6b,c), and include promoter shape, TATA-box, and the number of TFs binding to the promoter. An additional feature highly predictive of genes with low expression variation was the presence of CpG islands, in line with previous findings in single-cells (Morgan and Marioni 2018), while bivalent TSS state was predictive of high expression variation, in line with previous studies (Faure, Schmiedel, and Lehner 2017) (Fig. 6b, c). We also uncovered a number of transcription factors predictive of low variation, including GABPA, YY1, and E2F1 (84 predictive TFs in total, Supplementary Table 17). Similar to *Drosophila,* the presence of TSS-proximal peaks of all 84 predictive TFs were associated with reduced mean expression variation, again suggesting that high variation (in bulk RNA-seq) is due to a lack of buffering mechanisms rather than a specific mechanism for noise amplification. Extending the distance around the TSS did not improve the correlation between presence of TF peaks and expression variation, indicating that the key regulatory information is already contained within the core promoter region (Supplementary fig 8b).

We next asked whether expression variation across individuals is predictive of differential expression in different conditions, as we observed in *Drosophila*. For this we used differential expression prior (DE prior), a metric that integrates more than 600 published differential expression datasets and reflects the probability of a gene to be DE irrespective of the biological condition tested (Crow et al. 2019). Indeed, DE prior is correlated with expression variation in all tissues (median Pearson correlation R= 0.50), while being uncorrelated with expression level (Supplementary Fig. 6). A model trained to predict the top-30% vs. bottom-30% most variable genes (based on the features predictive of mean expression variation) could predict DE prior with an AUC of 0.75 versus 0.85 when both training and testing are done on DE prior (Fig. 6d, Methods), and predictive features for variation showed similar effects in DE prior (Fig. 6e). This indicates that inherent promoter features can explain expression variation and the probability of differential expression to a similar extent – potentially, due to partially overlapping underlying mechanisms.

Importantly, both expression variation and DE prior were significantly lower for essential genes, while being higher for GWAS hits and common drug targets (Fig. 6f, Supplementary Fig. 8c). Higher expression variation of the latter agrees with an interpretation that these genes are less buffered to withstand different sources of variation (Fig 1a) and hence are more likely to change in expression level upon different types of perturbations including genetic or environmental factors. Hence, expression variation across individuals likely captures differences in selection pressure and cost-benefit trade-offs between expression precision and plasticity.

In summary, despite significant differences in promoter regions between humans and *Drosophila* (e.g. the presence of *Drosophila*-specific core promoter motifs, human-specific CpG islands, predominately unidirectional versus bidirectional transcription), promoter features are highly predictive of expression variation in both species. Genes with high variation tend to also have differential expression across diverse conditions, and are significantly enriched in GWAS hits, and disease associated loci.

## Discussion

Our analysis suggests that expression variation across a population of multicellular genetically diverse individuals is gene-specific and can be explained by genetically encoded regulatory features, all highly correlated with core promoter architecture. Overall, we found that regulatory complexity positively correlates with robust gene expression. Yet we identified two independent mechanism that decrease expression variation depending on the core promoter architecture. Genes with broad core promoters in *Drosophila* were overall less variable and characterized by ubiquitously open chromatin and a high number of transcription factors (TFs) binding to the TSS-proximal region. In contrast, genes with a narrow core promoter had a much higher spread of expression variation, which was, in addition to TFs, modulated by regulatory complexity outside of core promoters (miRNAs, enhancers and Pol II pausing).

We found that similar promoter-related features were predictive of expression variation across human individuals by applying the same predictive framework to tissue-specific RNA-seq datasets. This was surprising given the differences in promoter features between *Drosophila* and mammals, with higher heterogeneity within broad promoters and high regulatory importance of CpG islands (Haberle and Stark 2018; Lenhard, Sandelin, and Carninci 2012), and suggests that some core promoter properties are ancient features that reduce expression noise, which agrees with conclusion of previous studies (Carey et al. 2013; Metzger et al. 2015).

Gene expression variation can arise from a multiplicity of stochastic, environmental and genetic factors, and defining the exact cause of expression variation in a particular experiment is likely an intractable task. Even for single cell experiments, which can control for genetic and macro-environmental factors, there is ongoing debate as to whether the observed gene-specific expression variation can be explained by intrinsic (e.g. transcription bursting) or extrinsic (cell-to-cell variability) factors (Battich, Stoeger, and Pelkmans 2015; Larsson et al. 2019; Foreman and Wollman 2019), or whether these are sources are indistinguishable (Eling, Morgan, and Marioni 2019). Yet, despite the differences in interpretation of the underlying sources of variation, there is a consensus that genes differ in their expression variation. Here, we found that gene expression variation, in bulk data from thousands of cells, was highly reproducible across different datasets, including developmental time-points in *Drosophila* and tissues in human, and did not depend on the identity of samples used. This suggests that gene expression variation, along with expression level, can be used as an informative readout of gene function and regulation in multiple biological contexts.

Interestingly, we recapitulated most of the regulatory features previously linked to expression noise in single cell experiments (Ravarani et al. 2015; Morgan and Marioni 2018; Faure, Schmiedel, and Lehner 2017; Perry et al. 2010; Boettiger and Levine 2009; Schmiedel et al. 2018), despite the fact that the composition of variation sources is very different between bulk and single cell experiments. A number of studies have proposed that robustness to stochastic noise and robustness to environmental and genetic variation are highly correlated (Lehner 2008; Ciliberti et al. 2007; Kaneko 2011). In line with this hypothesis, expression variation in bulk is predictive of single-cell noise in yeast (Dong et al. 2011) and gene expression variation across individuals in human tissue samples correlates with promoter strength and multiple epigenetic features (Alemu et al. 2014). Indeed, genes that have evolved mechanisms to buffer stochastic variation in the levels of their expression may also be insensitive to non-stochastic changes, including genetic and environmental variation, as the same mechanisms would constrain them both (Lehner 2008).

In line with the above, it was recently shown that the results of many differential expression experiments are generally predictable and to a large extent reflect some basic underlying biology of the genes, rather than specific conditions tested (Crow et al. 2019). Our results confirm and substantially extend this model - we show that the likelihood of a gene to be differentially expressed is highly correlated with the gene’s expression variation (independent of expression level) and the corresponding predictive regulatory features. This result is important, as standard differential expression pipelines correct for variance dependence on the expression level (Love, Anders, and Huber 2014) but do not take any other gene-specific properties into account. Given the extensive amount of accumulated knowledge about regulatory features, taking into account gene-specific differences in expression variation and the underlying regulatory mechanisms will improve specificity and interpretability of differential expression results.

Finally, here we focused on the most general mechanisms robustly linked to gene expression variation regardless of the specific tissue identity or developmental stage. There is, however, accumulating evidence that changes in expression variation can be an important indicator of specific biological processes happening in an organism. In particular, stochasticity of expression can differ by developmental stage i.e. following an hourglass pattern in early development (Liu et al. 2019) or decreasing with cell fate commitment (Eling et al. 2018; Richard et al. 2016). On the other hand, an increase in expression stochasticity has been linked to ageing (Viñuela et al. 2018; Kedlian, Melike Donertas, and Thornton 2019) and certain disease conditions (Zhang et al. 2015; Ran and Daye 2017). Hence, combining information on expected gene expression variation with tissue or disease-specific data might provide additional insights to condition-specific gene regulation in complex biological systems.

## Methods

### Gene expression level and variation in *Drosophila* DGRP lines

#### Gene expression quantification

To quantify gene expression, we re-processed the single-end strand-specific 3’-Tag-seq data (Cannavò et al. 2016) for 75 inbred wild *Drosophila* isolates from the *Drosophila melanogaster* Genetic Reference Panel (Mackay et al. 2012) at three time-points during embryonic development (2-4, 6-8 and 10-12 hours after fertilization, 225 samples in total, each containing pool of approximately 100 embryos). Reads were trimmed using Trimmomatic v.0.33 software (Bolger, Lohse, and Usadel 2014) with the following parameters: -phred33 HEADCROP:7 CROP:43. Alignment to dm6 genome version was done with bwa v.0.7.17 aln (parameters: -n 5 -e 10 -q 20) and samse (parameters: -n 1) tools (Heng Li and Durbin 2010). Reads with mapping quality below 20 were removed using samtools view v1.9 (H. Li et al. 2009). Expression was quantified with HTSeq count v.0.9.1 (Anders, Pyl, and Huber 2015) (parameters: -m intersection-nonempty -f bam -s yes -q -i Parent). PolyA sites were identified by reproducing the analysis of the polyadenylation dataset published in (Cannavò et al. 2016) after mapping the reads to the dm6 genome assembly. We observed a partial failure of strand specificity in generating the sequencing libraries: highly expressed polyA sites showed a corresponding antisense site. To remove these artefacts, we excluded polyA sites that were perfectly included in an antisense site. Reads that spanned both the last transcribed base and the subsequent polyadenylation tail allowed for single base resolution identification of the cleavage site. We extended polyA sites 200bp downstream or up to the nearest polyA site. To identify cleavage sites within our polyA sites we produced strand specific coverage tracks of the 3’-terminal base for each of the polyadenylation reads. Within each pA region, we identified the major cleavage site as the genomic base with highest 3’-terminal base coverage.

#### Expression data filtering and measuring expression variation

All samples selected for the analysis had high sequencing quality and were accurately staged, as described in original publication (Cannavò et al. 2016). Using principal component analysis on the expression counts from all 225 samples after applying variance stabilization transformation from DESeq2 (Anders and Huber 2010), we confirmed that samples clustered by developmental time-point (Supplementary fig 1a) and not sequencing batch (Suppl. fig 1b).

Expression counts from 225 samples were jointly normalized using effectSize normalization from DESeq2 package (Anders and Huber 2010). For each time-point separately, we calculated median expression and coefficient of variation (CV, standard deviation divided by mean) for each gene across 75 samples. Genes with zero median expression were removed as non-expressed. Coefficient of variation exhibited strong negative relationship with median expression level (Fig. 1b) which agrees with other gene expression studies (Anders and Huber 2010; Ran and Daye 2017; Faure, Schmiedel, and Lehner 2017; Eling et al. 2018). To account for this relationship, we used Locally weighed regression (LOESS) of coefficient of variation on the median expression (loess function in R from stats library, degree = 1, span = 0.75) (R Development Core Team 2013). Residuals from LOESS regression (resid_cv, residual coefficient of variation) were used in all subsequent analysis and referred to as gene *expression variation*.

To check whether residual expression variation actually reflects expression heterogeneity (across samples) at any given expression level, we took the following approach. Genes were grouped into 20 bins by their median expression level across 75 samples (separately for each time-point). Within each bin, genes were ordered by their residual coefficient of variation (x-axis), and normalized expression counts for each sample were plotted on the y-axis (example for 10-12h in Supplementary Fig. 1c). For almost all of the expression bins, spread of expression values increased for higher residual coefficient of variation, except bin-20 (top-5% genes by expression level) and to less extend bin-1 (bottom-5%). Based on this analysis, top and bottom-5% of expressed genes were excluded from the analysis.

We focused our analysis on the latest developmental stage (10-12h) and removed genes that decreased in expression between 2-4h and 10-12h after fertilization. This was done to reduce confounding effects of maternal mRNA degradation and focus on the stage when zygotic genome is fully activated (both processes happening from 2h post fertilization onwards). In total, we excluded 3275 genes, from which 90% were detected as maternally deposited (in house data, genes expressed in unfertilized eggs). In addition, genes with the strongest decrease in expression (3-fold or more) were highly enriched in cell cycle biological processes (Supplementary table 7), and cell cycle is known to slow down at later developmental stages (Edgar and O’Farrel 1989). Hence, we reasoned that variation of these genes might be strongly confounded by extrinsic factors (maternal mRNA degradation and cell cycle) that are not of particular interest for this analysis.

Overall, the following filtering steps were applied to the data, and the corresponding genes were excluded from the final dataset:

1. Genes with zero median expression level across samples (as non-expressed genes);
2. Genes falling into top and bottom 5% by expression level (as potential source of outliers);
3. Genes that decreased in expression between 10-12 and 2-4 hours after fertilization (as maternal genes with role in early embryonic development and potential targets for maternal mRNA degradation)
4. Genes with missing values in the feature table (see below) unless the feature can be easily imputed, i.e. 0 for the absence of transcription factor motif

Hence, our final dataset included 4074 genes at 10-12 hours post fertilisation. Final measure of expression variation was calculated as described above on the final set of genes to avoid residual dependence on the expression level after filtering (Fig. 1b, *‘resid_cv’* column in Supplementary table 3). Full dataset for all there time-points including expression variation calculated at several intermediate filtering steps are provided in Supplementary Table 2.

#### Expression variation on the subsets of samples

To test robustness of expression variation to the selection of samples (and hence potential batch effects), we performed multiple rounds of sample subsetting. Our full dataset comprised 75 samples (75 DGRP lines). For a given subset size N, we randomly selected N samples from the full dataset. Gene expression variation was calculated on this subset as described above (including fitting LOESS regression), and Pearson correlation of resulting variation values with the variation on the full dataset was recorded. Radom selection of samples was performed 100 times for each N. This was done for the following subset sizes: 5, 10, 20, 30, 40, 50, 60, 70, and 74 samples. Mean and standard deviation of correlation values upon 100 rounds of sampling for each subset size are shown in Fig 1c.

#### Expression level and variation of neighboring genes

For this analysis we considered all pairs of genes located on the same chromosomes and with TSS to TSS distance below 100 kB. Genes pairs were binned into 5 quantiles by the distance between their TSSs. Coordinates of the topologically associated domains (TADs) were taken from the high-resolution HiC in Kc cells (Ramírez et al. 2018). Genes were assigned to TADs based on their TSS coordinates, and for all pairs of genes we defined whether they belong to the same TAD or span the TAD border. Within the resulting 10 groups of gene pairs (5 quantiles * same/different TAD), we calculated Spearman correlation coefficients in expression variation and median expression level between genes in the pairs (Supplementary Fig 1d).

#### Alternative measures of expression variation

As alternative measures of expression variation, we used the following metrics:

1. sd_vst: standard deviation after applying variance stabilizing transformation from DESeq2 package to remove mean-variance dependence (instead of taking LOESS residuals)
2. resid_sd: LOESS residuals from regressing standard deviation on median expression
3. resid_mad: LOESS residuals from regressing median absolute deviation on median expression
4. resid_iqr: LOESS residuals from regressing interquartile range (between 25^th^ and 75^th^ percentiles) on median expression

These measures were calculated on the final set of 4074 genes at 10-12h post fertilization. Dependences on the median expression before and after correction for these measures are provided in Supplementary fig 2a. Pearson correlations with expression variation measured by resid_cv are shown in Suppl. fig 2b.

### Compiling Feature table for *Drosophila* dataset

Full list of features used in this analysis is provided in Supplementary table 1. The features were grouped into seven classes (column *‘Feature class’* in Supplementary table 1): Genetics, Gene type, Gene body, TSS, 3’UTR, Distal regulators, and Gene context. Summary on assignment of features to classes is provided in Table 1. Below are the more detailed descriptions of how individual features were generated. Full feature table and final dataset are provided in Supplementary tables 2 and 3, respectively.

#### Basic gene properties and functional annotations

We used Flybase v6.13 genome annotation to find gene length (length_nt), number of transcripts (n_transcripts) and number of exons (n_exons) for each gene. Number of exons was defined as total number of unique exons regardless of transcript isoforms. Next, we used several gene annotations from in-house or external sources to identify specific functional groups of genes. Ubiquitously expressed genes (is_ubiquitous) were defined based on BDGP database (Tomancak et al. 2002) as genes having ubiquitous expression pattern in at least one developmental stage (data available for *Drosophila* embryonic stages 4-6, 7-8, 9-10, 11-12, 13-16). Maternally deposited genes (is_maternal) were defined as genes expressed in unfertilized eggs the vgn line of *Drosophila melanogaster* at 2-4 or 6-8 hours after egg laying (in-house data, unpublished). Housekeeping genes (is_housekeeping) were defined following methodology in (Ulianov at al. 2015) as genes expressed with RPKM > 1 in all samples from (Graveley et al. 2010). List of transcription factors (is_tf) comes from (Hammonds et al. 2013) dataset.

#### Human orthologs for Drosophila genes

Human orthologs for *Drosophila melanogaster* genes were identified with DIOPT - DRSC Integrative Ortholog Prediction Tool (Hu et al. 2011), and two features provided by the tool were added for each gene – conservation score (conserv_score, continuous variable indicating confidence of ortholog prediction) and conservation rank (conserv_rank, factor variable taking the following values: none, low, moderate, high). Genes with ‘high’ conservation rank were referred to as “conserved with human” (e.g. Supplementary Fig 3c).

#### Genetics

*Cis share* (cis) was used as an estimate of the contribution of genetic variation to the total gene expression variation. To calculate it, we used LIMIX variance decomposition (Lippert et al. 2014) on normalized expression matrix (three time-points combined) to assess the proportion of gene expression variation explained by cis, population structure and time/environment. *3’UTR variant index* (utr3_variant_index) was used to approximate a potential effect of mappability bias (because expression was estimated from 3’Tag-seq data) as well as sequence variation on gene expression variation. It was calculated with the following formula: (total number of variants in gene’s 3’UTR × mean allele frequency of variants) / total length of 3’UTR peaks. The variant counts and variant allele frequencies were obtained from the DGRP freeze 2 .vcf file (W. Huang et al. 2014), considering only the 75 lines used in this study. *Presence of eQTL* (with_eQTL) indicates whether a gene has associated expression QTL identified in (Cannavò et al. 2016) on the expression dataset, which is also used in this study.

#### GC-content

GC-content was calculated using bedtools-2.27.1 nuc software (Quinlan and Hall 2010) for nucleotide sequences of genes (gene_gc) and regions of −100/+50 bp around gene TSS annotations from Flybase v6.13 (tss_gc).

#### Pausing Index, Promoter shape and promoter motifs

Polymerase II pausing index (defined as the density of polymerases in the promoter region divided by the gene body) in Drosophila melanogaster embryos was taken from (Saunders et al. 2013).

Promoter shape index was defined in the earlier paper (Schor *et al*., 2017) following the methodology from (Hoskins et al. 2011). In brief, promoter shape index is Shannon entropy of the TSS distribution within a promoter:

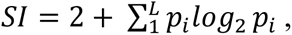

where p is the probability of observing a TSS at base position i within the promoter, L is the set of base positions that have at least one TSS tag, and TSS positions were identified using the aggregated CAGE signal for all time points and 81 fly lines from the Drosophila Genetic Reference Panel (DGRP) at three developmental time-points (Schor et al. 2017). For each gene, we recorded promoter shape index of the most expressed TSS cluster (major_shape_ind). Promoters of genes were classified as broad if shape index of the most expressed TSS was below −1, and narrow otherwise. The threshold is based on the bimodality of shape index distribution and was defined in the original publication (Hoskins et al. 2011). If any of alternative TSSs of a gene had shape different from the most expressed one, alt_shape feature took value of 1 (and 0 otherwise).

PWMs for 8 core promoter motifs (Ohler et al. 2002; Ohler 2006) were scanned in −100/+50 bp region around annotated TSSs from Flybase v6.13 using fimo-4.11.3 software (Bailey et al. 2009) with uniform background (--bgfile –uniform--), no reverse compliment (–norc), and default p-value threshold (1e-4). Motifs were first scanned for the 5’-most TSS of each gene (start coordinate of genes in gff annotation) and referred to as ‘ohler_maj.motif_name’ (e.g. ohler_maj.TATA for TATA-box; 0/1 for motif absence/presence respectively). In addition, motifs were scanned for TSSs of all transcripts for each gene (start coordinates of transcripts in gff annotation). If motif was predicted for some of the transcript TSSs but not for the gene TSS, then the corresponding feature ohler_alt.motif_name took value of 1, otherwise 0.

#### DNase hypersensitive sites

DNase hypersensitive sites (DHS) in *Drosophila melanogaster* embryos were identified in [Reddington et al, submitted]. The experiment was conducted at four developmental time-points in whole embryo (2-4h, 4-6h, 6-8h, and 10-12h after fertilization) and with tissue sorting (mesoderm, neuroectoderm, and other (double negative) at all time-points except 2-4h; bin-positive and bin-negative mesoderm (marker for visceral muscles) at 6-8h). This resulted in 19 experiments, which we refer to here as *DHS conditions*. Peaks called in all experiments were combined in a single table, and for each DHS, conditions when the site was accessible were recorded. Coordinates of DHS peaks from the combined table were lifted over from dm3 to dm6 genome version using UCSC liftOver-5.2013 tool (Kent et al. 2002).

For each gene, we quantified a number of features related to DHS in TSS-proximal (+/-500 bp. around TSS from gene annotation, class TSS) or TSS-distal (more than 500 bp and less than 10kB around annotated TSS, class Distal regulators):

- Number of conditions with DHS (num_dhs_conditins.prox and num_dhs_conditions.dist) is the number of conditions (out of 19 in total) when there was a DHS peak detected in TSS-proximal or TSS distal region.
- DHS tissue profile (dht_tissue_profile.prox and dhs_tissue_profile.dist) summarizes accessibility profile across tissues and takes the following values: 1 – peak present only in tissues (any of mesoderm, neuroectoderm and double negative); 2 - present in whole embryo (WE); 3 – both in WE and tissues.
- DHS time profile (dhs_time_profile.prox and dhs_time_profile.prox) reflects accessibility profile across developmental time points: 1 – peak present only at early developmental time-points (2-4h, 4-6h or 6-8h after fertilization); 2 – peak present only at late developmental time-points (8-10h or 10-12h after fertilization); 3 – peak present in at least one early and late time-point.
- Presence of ubiquitous DHS (is_ubiq.prox and is_ubiq_dist) indicates presence of ubiquitously accessible DHS peak in the corresponding genomic region. We consider DHS peak ubiquitous if it was present in all three tissues at four developmental time-points where tissue sorting was done (12 conditions in total).
- Number of DHS peaks (num_dhs_any.prox and num_dhs_any.dist) is the total number of non-overlapping DHS peaks in the corresponding intervals present in any of the 19 conditions.

#### DNA binding proteins

280 embryonic ChIP-seq datasets for various DNA binding proteins (referred to as transcription factors or TFs for simplicity though not all of them have transcription factor activity) were downloaded from modERN database (Kudron et al. 2018). Of note, for several transcription factors, ChIP-seq data are available either for several developmental time-points (Trl at 0-24h, 8-16, and 16-24h. after fertilization) or for several experimental setups (chif-RA-GFP and chif-RB-GFP). In case more than one data set was available for a TF it was included independently. For the analysis, we used peaks called according to the methodology from the original publication (IDR threshold of 0.01, optimal set). ChIP-seq peaks were overlapped with DHS coordinates (single base pair overlap required) using findOverlaps function from GenomicRanges package in R (Lawrence et al. 2013) resulting in 280 binary variables (1/0 for presence/absence of each TF) were added to the DHS table. These data were then summarized for each gene’s TSS-proximal and TSS-distal region resulting in the following variables:

- Presence of TF peak in TSS-proximal DHSs (280 variables with name format like modERN.tf_name.prox) and TSS-distal DHSs (280 variables with name format like modERN.tf_name.dist); 1 – peak present (any number of occurencies), 0 – peak absent.
- Total number of different TF peaks overlapping TSS-proximal (num_tf_peaks.prox) and TSS-distal (num_tf_praks.dist) DHS.

640 PWMs for different TF binding motifs (Drosophila melanogaster database, version available on 05.03.2019) were downloaded from CIS-BP database (Weirauch et al. 2015). PWMs were scanned in the sequences of DHSs resized to 200 bp. width using fimo-4.11.3 software (Bailey et al. 2009) with uniform background (--bgfile –uniform--), with reverse compliment (default), and default p-value threshold (1e-4). Similar to TF peaks, these data were then summarized for each gene’s TSS-proximal and TSS-distal region resulting in the following variables:

- Presence of TF motif in TSS-proximal DHSs (280 variables with name format like cisbp.tf_name.prox) and TSS-distal DHSs (280 variables with name format like cisbp.tf_name.dist); 1 – motif present (any number of occurencies), 0 – motif absent.
- Total number of different TF motifs overlapping TSS-proximal (num_tf_motifs.prox) and TSS-distal (num_tf_motifs.dist) DHS.

#### Chromatin colours

Annotation of chromatin states (5 states) was taken from (Filion et al. 2010). Coordinates of genomic regions assigned to different colours were overlapped with DHS table, and for each DHS overlap with any of the colours by at least 1 bp. was recorded. The results were aggregated by gene into 5 TSS-proximal (i.e. color_green.prox) and 5 TSS-distal (i.e. color_green.dist) binary features indicating presence/absence of the corresponding states.

#### Annotated enhancers

We used several datasets of annotated enhancers from the following sources:

- Combined set of CAD4 enhancers (curated in-house list from various sources) and Vienna tiles (Kvon et al. 2014) lifted over to dm6 genome version;
- Combined set of cis-regulatory modules (CRMs) of mesoderm TFs (Zinzen et al. 2009) and cardiac TFs (Junion et al. 2012) lifted over to dm6 genome version;

Both datasets were first overlapped with DHS table and number of annotated enhancer elements in TSS-proximal and TSS-distal regions were added to the feature table.

#### 3’UTR features

PWM of micro-RNAs (miRNAs) from MIRBASE (Kozomara and Griffiths-Jones 2014; Kozomara, Birgaoanu, and Griffiths-Jones 2019) and RNA-binding proteins (RBPs) from CISBP-RNA (Ray et al. 2013) were downloaded from MEME v4 (Bailey et al. 2009), files *Drosophila_melanogaster*_dme.dna_encoded.meme and *Drosophila_melanogaster*.dna_encoded.meme for miRNA and RBP PWMs respectively. 3’UTRs were defined as the region comprised between a major cleavage site (as defined above) and the closest annotated stop codon. PWMs were scanned in nucleotide sequences of the 3’UTRs using fimo-4.11.3 software (Bailey et al. 2009) with uniform background (--bgfile – uniform--), no reverse compliment (–norc), and default p-value threshold (1e-4). Features for motif occurrences have were named mirbase.motif_name and cisbp_rna.motif_name for miRNA and RBP motifs respectively. The feature took value of 1 for a gene if the corresponding motif was predicted for any of the annotated 3’UTRs of a gene and 0 otherwise. Total number of unique miRNA and RBP motifs per gene were counted and included as num_mirna and num_rbp features respectively.

Lists of genes that are putative targets of Pumilio (embryonic and adult data) and Smaug (embryonic data) RBPs were obtained from (Gerber et al. 2006) and (Chen et al. 2014), respectively.

For each gene, we calculated the mean UTR length at different time points as the weighted mean UTR length between UTR isoforms. We used the polyA site expression as weights in the mean calculation. Since length of 3’UTR was highly correlated with gene length (Spearman correlation, Rho=0.62), utr3_length feature was calculated as actual 3’UTR length divided by gene length. Finally, 3’UTR length changes (log2-fold change) between different time-points (10-12h vs. 6-8h, 6-8h vs. 2-4h, 10-12h vs 2-4h) were calculated for each gene (utr3_l2fc_10vs6, utr3_l2fc_6vs2, and utr3_l2fc_10vs2 features).

#### Genomic context features

Insulation score (ins_score_2_4h and ins_score_6_8h) was calculated based on Hi-C data in-house data (unpublished) for *Drosophila melanogaster* embryos at 2-4 and 6-8 hours after fertilization (in-nucleus ligation, whole embryo). To assign insulation score to genes, we recorded the nearest value to the annotated TSS of each gene.

Coordinates of topologically associated domains (TADs) were taken from the high-resolution Hi-C in Kc cells from (Ramírez et al. 2018) and Hi-C in 2-4h embryos (in-house data, unpublished). Each gene was then assigned to TAD from the two aforementioned annotations based on its TSS coordinate, and distance to TAD border and TAD size were recorded (dist_to_tad_border.ramirez, dist_to_tad_border.2_4h, tad_size.ramirez, and tad_size.2_4h, respectively).

Gene density was calculated as number of genes in +/-1000 bp and +/-20kB from the TSS of each gene (num_genes.prox and num_genes_dist, respectively) based on Flybase v6.13 genome annotation.

#### Broad and narrow indices

Broad and narrow indices were calculated based on the subset of features from the feature table. Broad index was composed of the following features (all TSS- proximal): number of conditions with DHS (num_dhs_conditions.prox), number of TF peaks (num_tf_peaks.prox), number of TF motifs (num_tf_motifs.prox). Narrow index was composed of number of TSS-distal DHSs (num_dhs_any.dist), number of miRNA motifs (num_miRNA), and Pol II pausing indes (PI). All features were first converted to ranks (random order for ties). Indices were calculated as simple averages of the corresponding features.

### Measuring expression level and variation in human tissues

#### Genome version

We used Ensembl GRCH37/hg19 genome version downloaded from UCSC table browser (Kent et al. 2002; Haeussler et al. 2019) throughout the analysis. Sex chromosomes and non-standard chromosomes were removed for all subsequent analyses. For selecting the main transcript per gene we used GRCH37/hg19 genome annotation downloaded from Ensembl website (Cunningham et al. 2019).

#### Quantifying expression level and variation

Gene expression matrix (raw read counts) was downloaded from the GTEx project website (GTEx Consortium 2013). Gene read counts matrices per tissues were produced by using GTEx sample details downloaded from GTEx website. Tissues with more than 100 samples (43 tissues in total) were chosen for further analysis (Supplementary table 8). In each tissue, genes with 0 median counts were removed and expression counts were normalized using size factor normalization form DESeq2 package in R (Love, Anders, and Huber 2014). Median expression levels were calculation for each gene in each tissue and converted to log-scale (natural logarithm) for subsequent analysis.

Next, we removed top-5% of genes by median expression level as potential outliers, following the same reasoning as for Drosophila. Since distributions of gene expression in all tissues had long left tails, we set additional stringent threshold on lowly expressed genes (minimum median of 5).

Gene expression variation was calculated on the final set of genes for each tissue following the same approach as for Drosophila. Namely, gene expression variation was defined as the residuals from the local linear regression of coefficient of variation (CV) on the median expression (both on the log-scale, loess function in R from stats library, span = 0.25 and degree = 1). Gene expression levels and variations in all tissues are provided in Supplementary table 9.

Mean expression variation for each gene was calculated as the mean of expression variations in all tissues where a gene was expressed using final tables that passed all filtering steps. Similarly, mean expression level was calculated by computing the mean of median expression levels in all tissues where a gene was expressed. Mean expression variation calculated in this way exhibited weak dependence on mean expression level (Spearman correlation, Rho=-0.11). To control for this effect, we also calculated ‘global mean variation’ as the residuals from the local linear regression of the mean CV on the mean expression level (calculated as above). This measure was highly correlated with mean variation (Supplementary fig S6) and showed similar results in the downstream analysis (results not shown, global mean variation is provided in Supplementary Table 9).

### Feature tables for human dataset

Only TSS-proximal features and several gene properties (i.e. gene length and number of transcripts) were used to predict expression level and variation in human. Full list of features used in this analysis is provided in Supplementary table 10. Most of the TSS-proximal features (TF peaks and chromatin states) were scanned in the −500/+500 bp of the main TSS of the genes (referred to as *TSS-proximal regions*), following the same approach as for *Drosophila*. Several features more strictly linked to the gene core promoters (promoter shape, TATA-box, CpG islands, and promoter GC-content) were scanned in −300/+200 bp of the main TSS of the genes (referred to as *core promoter regions*).

#### Gene properties

Number of transcripts, gene length, mean exon length, number of exons and exon length mean absolute deviation were calculated for each gene directly using hg19 genome annotation from Ensembl website (Cunningham et al. 2019). Transcripts width was calculated for each transcript by using the same file, and length of the main transcript was assigned to each gene.

#### Promoter Shape

CAGE data for 31 tissues (library size of about 10M mapped reads or above, Supplementary table 11) was downloaded from FANTOM5 project (Lizio et al. 2015) using CAGEr package in R (Haberle et al. 2015). On each dataset separately, we did power-law normalization (Balwierz et al. 2009) using CAGEr package. TSSs with low count numbers (less than 5 counts) were removed. Next, we applied a simple clustering method (distclu, maximum distance = 20) form CAGEr package on each dataset separately. Clusters with low normalized CAGE signals (sum of TSSs normalized signals of the cluster below 10-50 depending on the tissue) were removed. CAGE clusters were then assigned to genes by overlapping them with core promoter regions (−300/+200 bp around TSSs of all annotated transcripts). Clusters that did not overlap any core promoters were removed.

Next we defined promoter shape for all CAGE clusters by using two commonly accepted metrics: promoter width and promoter shape index. Promoter width was calculated by using the inter-percentile width of 0.05 and 0.95 following methodology from (Haberle et al. 2015). Promoter shape index (SI) was calculated by the formula as above (*Drosophila* section) proposed in (Hoskins et al. 2011).

For classifying promoters into broad and narrow based on promoter width, we used the following approach. First, we did a linear transformation of promoter width values (actual value minus 1 divided by 10; for fitting gamma distribution) On the transformed data, we fitted gamma mixture model (2 gamma distribution), and parameters was trained using EM algorithm (Dempster, Laird, and Rubin 1979) using mixtools package in R (Benaglia et al. 2009). The threshold for classifying promoters as broad or narrow was selected by finding the point which best separates the two distributions. Following this approach, promoters with width above about 10-15 bp. were classified as broad, which was consistent across all tissues and agreed with earlier studies (Forrest et al. 2014). To classify promoters into broad/narrow using shape index, we fitted Gaussian mixture model (2 Gaussian distribution) to the data and selected the threshold separating the two distributions using the same approach as above. For the subsequent analysis, we used promoter width feature since it showed more clear bi-modal distribution in all tissues (example in Supplementary fig 7a-c) and is a more common metrics in the analysis of mammalian promoters (Forrest et al. 2014; Carninci et al. 2006).

Each gene was then assigned the promoter width of its main transcript. If more than one CAGE cluster was present for a gene’s main transcript, the cluster with the highest normalized CAGE signal was selected. Promoter width values for most of the genes were highly correlated across tissues (Supplemetary fig 7d). Based on the tissue-specific shape data, we calculated two aggregated features for each gene. Mean promoter width (mean_width feature) was calculated as the mean of gene promoter widths in all tissues where it had CAGE signal (passing the filtering criteria defined above). Share of tissues where a gene had broad promoter (percentage_of_broad feature) was calculated for each gene by dividing the number of tissues where the gene had broad promoter by the total number of tissues where the gene had CAGE signal.

#### TATA-box motif

TATA-box motif coordinates were obtained from the PWMTools web server (Ambrosini, Groux, and Bucher 2018): JASPAR core 2018 vertebrates motif library (Khan et al. 2018), p-value cutoff of 10-4, GRCh37/hg19 genome assembly). Motif coordinates were overlapped with gene core promoter regions (−300/+200 bp), and number of overlaps for each gene was recorded (TATA_box feature).

#### Transcription Factors

Transcription factors dataset (444 TFs, peaks with motifs, hg19 genome) ware obtained from (Vorontsov et al. 2018). If several datasets were available for the same TF, the dataset with the best quality was selected. For each TF, the corresponding feature was calculated by overlapping the TF regions and gene TSS-proximal regions (−500/+500 bp) and counting the number of overlaps for each gene.

#### Chromatin States

Chromatin States dataset (chromHMM core 15-state model with 5 marks and 127 epigenomes (Ernst and Kellis 2017)) was downloaded from Epigenomics Roadmap project (https://egg2.wustl.edu/roadmap/web_portal/chr_state_learning.html). We considered 26 tissues (Supplementary table 12). For each tissue, 15 features (one for each state, e.g. TssA or TssBiv) were obtained. Each feature was calculated by overlapping corresponding state regions and gene TSS-proximal regions (−500/+500 bp) and counting the number of overlaps for each region. Finally, aggregated features (e.g. mean_TssA or mean_TssBiv) were calculated as the mean of feature values for each state over all 26 tissues.

#### CpG Islands

CpG islands (CGI) data for hg19 were downloaded from the UCSC Genome browser (Haeussler et al. 2019). For each CGI, these included CGI length (CpG_Length), number of CpG clusters (CpGNum) and number of GC dinucleotides (gcNum). The three corresponding features for each gene were calculated by overlapping CGI regions and gene core promoter regions (−300/+200 bp). When a gene did not overlap any CGI, the three features were assigned to 0. If multiple overlaps were present, CGI with the biggest overlap was considered for each gene.

#### Promoter GC-Content

Promoter GC-content (GC_content) was calculated by using biostring package (Pagès H et al 2019) and BSgenome.Hsapiens.UCSC.hg19 v1.4.0 in R in gene core promoter regions (−300/+200 bp).

#### Compiling final feature tables

We collated three tissue-specific datasets for lung, muscle, and ovary by combining the above promoter features and tissue-specific expression data (Supplementary tables 13-15). These tables included three types of features:

- Tissue-specific features (promoter width and chromatin states);
- Features aggregated across tissues (mean promoter width, percentage of broad, mean chromatin states – see above);
- Non tissue-specific features (all other features, e.g. TATA-box or TF peaks)

These tables included genes that were expressed and had CAGE signal (passing the above filtering criteria in both datasets) in the corresponding tissues. For muscle tissue, ‘Skeletal muscle male’ dataset was used for tissue-specific chromatin states. The fourth feature table included only non-tissue-specific and aggregated features along with mean expression level and variation (Supplementary table 16). This table was comprised of genes that were expressed and had CAGE signal in a least one of the analysed tissues. Expression variation was adjusted for the expression level on these final sets of genes in each table (see above).

#### Essential genes, drug targets, and GWAS catalogue

Essential genes (essential in multiple cultured cell lines based on CRISPR/Cas screens (Hart et al. 2017)) and drug targets (FDA-approved drug targets (Wishart et al. 2018) and drug targets according from (Nelson et al. 2015)) were downloaded from Macarthur lab repository (https://github.com/macarthur-lab/gene_lists). GWAS dataset was downloaded from EBI GWAS catalog (Buniello et al. 2019). Genes with GWAS associations within upstream regions or downstream regions were considered. These gene annotations were used in Fig. 6f and Supplementary fig. 8c, but not included in prediction models. Information on these gene types is provided in Supplementary table 9.

### Predicting expression level and variation

#### Random forest models for Drosophila embryos

Feature selection was done with the Boruta algorithm implemented in R (Kursa and Rudnicki 2010) with the following parameters: p-value = 0.01, maxRuns = 500; Z-scores of mean decrease accuracy measure as importance attribute; ranger implementation of random forest regression. Feature selection was done separately for several tasks: (1) predicting expression variation; (2) predicting expression level (log-transformed values); (3) predicting promoter shape index; (4-5) predicting expression variation and level in broad and narrow promoter genes separately. Median feature importance from 500 iterations were used as feature importance metrics. All features selected in at least one of the 5 setting listed above are provided in Supplementary table 5 with the corresponding importance. Only selected features were used in random forest predictions and all downstream analysis.

For each explained variable (expression variation, level or promoter shape index), we ran random forest regressions using mlr package in R (Bischl et al. 2016) with ranger implementation of random forest (Wright and Ziegler 2015); default parameters: num.trees = 500, mtry = square root of the number variables). Model performance was reported with coefficient of determination (R2) based on five-fold cross validation (Fig 1d).

#### Random forest models for human tissues

As above, we used random forest regression to predict expression level and variation in three tissues (lung, ovary, and muscle), as well as mean expression level and variation. Feature selection and random forest regression were performed in the same way and with the same parameters as for *Drosophila* dataset. Boruta feature selection algorithm was used to select important features predictive of expression level (log-transformed) and expression variation in each of the four datasets (three tissues and average). Feature importance scores are reported in Supplementary Table 17. Random forest regressions were run on the sets of selected features for the corresponding datasets. Model performance was reported with coefficient of determination (R2) based on five-fold cross validation (Fig 6a for performance in all 4 datasets).

### Testing robustness of the random forest models

#### Robustness tests for Drosophila dataset

We have run the following models to test robustness of our predictions to various potential confounding factors:

1. Binning genes by their median expression level into 5 quantiles and rerunning variation prediction for each quantile separately (Fig 1e);
2. Predicting alternative variation measures (see above): resid_sd, resid_mad, resid_iqr, and sd_vst (Supplementary Fig 2c);
3. Binning genes by their median expression change between 10-12 and 6-8 hours after fertilization into 5 quantiles and rerunning variation prediction for each quantile separately (Supplementary Fig 2d);
4. Binning genes by their promoter shape index into 4 quartiles and rerunning variation prediction for each quartile separately (Fig 3b).
5. Training and predicting on different chromosomes (or chromosome arms), e.g. leaving out all genes on chr3L for testing the model trained on all other genes (Supplementary Fig 2e).

For these tests, random forest regressions were run with the same parameters as above and on the set of features selected for the variation prediction on the full set of genes. Performance of the models measured with R^2 on the five-fold cross-validation in 1-4 and on holdout chromosome (arm) in 5.

#### Robustness tests on human datasets

Since human gene expression datasets from GTEx project contain high sample heterogeneity (different ages, sexes, reasons of death etc), we have rerun prediction models on the following subsets of individuals (using samples metadata from GTEx website) for the lung tissue expression dataset:

- Only 20-39 year old individuals;
- Only 40-59 year old individuals;
- Only 60-79 year old individuals;
- Only males;
- Only females
- Only violence group (as the reason of death);
- Only non-violence group (as the reason of death)

Gene expression variation and level were recalculated on the corresponding subsets of samples using the same methodology as above. Random forest regressions were rerun with the same parameters as above and on the set of features selected for the variation prediction on the full set of samples. Performance of the models was measured with R^2 on the five-fold cross-validation (Supplementary fig. 8a).

### Predicting differential expression in *Drosophila*

Lists of differentially expressed genes were obtained from (Moskalev et al. 2015). All experiments were conducted in adult *Drosophila melanogaster* flies (five-day old males) and included the following stress condition: entomopathogenic fungus infection (10 CFU, 10 CFU), ionizing radiation (144 Gly, 360 Gly, 864 Gly), starvation (16 h), and cold shock (+4°C, 0°C, −4°C). In total, 1356 out of our final set of 4074 genes were detected as differentially expressed in at least one of the above stress conditions (DE) versus 2718 non-DE genes.

To test how well model trained on expression variation can predict differential expression, we reformulated variation prediction into classification task to predict top-30% (class = 1) vs. bottom-30% (class = −1) of genes ranked by their expression variation (our embryonic dataset) and used trained model to predict DE (class = 1) versus non-DE genes (class = −1). To avoid having same genes in test and train sets, we undertook the following approach. Randomly sampled 50% of DE genes (678) and sample number of non-DE genes were set aside for train set. From the remaining genes (after excluding test set - either all 2718 genes, or only genes with narrow promoters), top-30% and bottom-30% of genes ranked by expression variation were used for training. Model was trained on the test set using random forest classification with default parameters (mlr package; ranger implementation of random forest; default parameters: num.trees = 500, mtry = square root of the number variables). Training was performed on the features important for predicting expression variation on the full set of genes (see above, Supplementary table 4) for expression variation (1 for high-variable, −1 for low-variable). Testing was done on the same set of features for differential expression (1 for DE, −1 for non-DE). Performance on the test set was assessed by Area Under the ROC curve (AUC). 10 rounds of random sampling of genes were performed, and mean AUC was reported (Fig 5d).

### Predicting differential expression prior in human

Differential expression Prior data (DE Prior rank) was obtained from (Crow et al. 2019). Ensembl ids were converted to entrez ids by using BioMart package in R (Durinck et al. 2009). We had information on both DE prior and mean variation (average of 43 tissue-specific variations across individuals, see above) for 11312 human genes. As above, we reformulated variation prediction into classification task to predict top-30% (class = 1) vs. bottom-30% (class = −1) of genes ranked by their expression variation and used trained model to predict top-30% (class = 1) versus bottom-30% (class = −1) genes ranked by DE prior. Training and testing were performed on the set of features predictive of mean expression variation in the main dataset (Supplementary table 17). Training and testing were done on the non-overlapping sets of genes using the following approach. First, we defined top-prior (top-30% by DE prior) and bottom-prior genes (bottom-30% by DE prior). 50% of genes from both groups were randomly sampled and assigned to test set. From the remaining genes, top-30% and bottom-30% by mean expression variation were selected for train set. The model was trained on the test set to classify top versus bottom variable genes (random forest classification with default parameters; mlr package in R, ranger implementation of random forest). Trained model was then used on the test set to predict top versus bottom DE prior genes. Similarly, another model was both trained and tested on classifying top versus bottom DE prior genes on the same train and test sets, respectively. Performance of the models on the test set was assessed by Area Under the ROC curve (AUC). 10 rounds of random sampling of genes were performed, and mean AUC was reported (Fig 6d).

### Statistical data analysis and visualization

Data analysis in R was done using base, stats, MASS, rcompanion, psych, tidyverse, magrittr, data.table, ltm, yaml, Boruta, mlr, ranger, GenomicRanges, DEseq2, CAGEr, and rtracklayer packages. All plots were done in R using ggplot2, ggpubr, gridExtra, ggExtra, RColorBrewer and pheatmap libraries. Contour lines in in Fig 2b-e represent 2D kernel density estimations (geom_density_2d with default parameters). P-values on the plots (Fig 2 b-e, Fig 4a-c,Fig 5b-c, Fig 6c,e,f) come from Wilcoxon rank test. Whiskers on the plots (Fig 1c-e, Fig 2f, Fig 3b, Fig 6a) indicate one standard deviation around the mean.

#### Correlation analysis

Generally, we used Spearman coefficient of correlation (R base) for comparing pairs of continuous variables or discrete variables taking more than two values (e.g. expression variation and promoter shape index or expression variation and conservation rank). In some cases, we used Spearman correlation coefficient (R base) to compare variables that are on the same scale, e.g. expression variations at different-time-points or for neighboring genes (same for comparing expression levels). Finally, point-biserial correlation coefficient (R, ltm library) was computed between continuous and binary variables (e.g. expression variation and presence of TATA-box motif). Median expression levels were log-transformed before computing correlation.

#### Gene Ontology enrichments

Gene Ontology (GO) enrichment tests were performed using clusterProfiler package in R (Yu et al. 2012). We used compareCluster function (p-value cut-off=0.01) to find enriched biological processes (Fig. 3c) and molecular functions (Supplementary fig. 3a) in genes grouped by their promoter shape and expression variation. For this analysis, genes with broad and narrow promoters were separately split into four quantiles by their expression variation (1-4 x-axis labels in Fig. 3c and Supplementary fig. 3a indicate quantiles: from low to high variation). Quantile intervals for broad promoter genes (1 to 4): [−1.06,−0.444]; (−0.444,−0.266]; (−0.266,−0.0754]; (−0.0754,1.89]. Quantile intervals for narrow promoter genes (1 to 4): [-0.98,-0.173]; (−0.173,0.0751]; (0.0751,0.416]; (0.416,1.99]. Full results of GO enrichment tests are provided in Supplementary tables 5 and 6.

#### Fisher’s tests

We used Fisher’s exact test (R base package) to find enrichments of features in different gene groups in *Drosophila* dataset (broad, narrow-low, narrow-high). All tests were done for 2×2 contingency tables, and odds ratios and p-values provided by the test were recorded. We used Benjamini-Hochberg correction to adjust p-values for the multiple testing. We used adjusted p-value threshold of 0.01 and odds ratio above 2 to define significantly enriched features (p-value adjusted < 0.01; odds ratio < 0.5 for significantly depleted).

First, we tested enrichment of housekeeping genes, transcription factors and TATA-box promoter motifs in the following pairwise comparisons: (1) broad vs. narrow, (2) narrow-low vs. two other groups, (3) harrow-high vs. two other groups. P-values were corrected for the number of tests (9 comparisons).

Next, we tested enrichments of ChIP-seq peaks of 24 transcription factors in the TSS-proximal regions in the same comparisons as above. 25 TSS-proximal TF features selected by Boruta algorithm, including two ChIP-seqs for Trl (in embryos at 8-16 and 16-24 hours after fertilization) from which the one with the overlapping time window was used (8-16h). Since ChIP-seq peaks were first overlapped with DHS peaks before assigning to genes (see Features section above), we restricted the analysis of TF enrichments to the genes that have at least one DHS peak in their TSS-proximal regions. P-values were corrected for 72 comparisons (24*2). Log2-transformed odds ratios from these tests are shown in Fig. 4c, weak enrichments/depletions (odds ratios above 0.5 and below 2) are shown in grey, actual values are provided in Supplementary table 5.

Peaks of Trl and Jarid2 (enriched in narrow-low) also showed weak enrichments in narrow-high, which likely comes from strong depletion of these TFs at the TSSs of broad promoter genes. To control for that, we also tested enrichments of the same 24 TF peaks in three comparisons between gene groups: (1) broad vs. narrow-low, (2) broad vs. narrow-high, (3) narrow-low vs. narrow high (also 72 comparisons for p-values correction).

Results from all Fisher’s tests described above are provided in Supplementary table 5.

## References

Alemu, Elfalem Y, Joseph W Carl Jr, Hé Ctor, Corrada Bravo, and Sridhar Hannenhalli. 2014. “Determinants of Expression Variability.” https://doi.org/10.1093/nar/gkt1364.

Ambrosini, Giovanna, Romain Groux, and Philipp Bucher. 2018. “PWMScan: A Fast Tool for Scanning Entire Genomes with a Position-Specific Weight Matrix.” Bioinformatics (Oxford, England) 34 (14): 2483–84. https://doi.org/10.1093/bioinformatics/bty127.

Anders, Simon, and Wolfgang Huber. 2010. “Differential Expression Analysis for Sequence Count Data.” Genome Biology 11 (10): R106. https://doi.org/10.1186/gb-2010-11-10-r106.

Anders, Simon, Paul Theodor Pyl, and Wolfgang Huber. 2015. “HTSeq-A Python Framework to Work with High-Throughput Sequencing Data.” Bioinformatics 31 (2): 166–69. https://doi.org/10.1093/bioinformatics/btu638.

Arnold, Cosmas D, Muhammad A Zabidi, Michaela Pagani, Martina Rath, Katharina Schernhuber, Tomáš Kazmar, and Alexander Stark. 2016. “Genome-Wide Assessment of Sequence-Intrinsic Enhancer Responsiveness at Single-Base-Pair Resolution.” Nature Biotechnology 35 (2): 136–44. https://doi.org/10.1038/nbt.3739.

Bailey, Timothy L, Mikael Boden, Fabian A Buske, Martin Frith, Charles E Grant, Luca Clementi, Jingyuan Ren, Wilfred W Li, and William S Noble. 2009. “MEME SUITE: Tools for Motif Discovery and Searching.” Nucleic Acids Research 37 (Web Server issue): W202–8. https://doi.org/10.1093/nar/gkp335.

Balwierz, Piotr J., Piero Carninci, Carsten O. Daub, Jun Kawai, Yoshihide Hayashizaki, Werner Van Belle, Christian Beisel, and Erik van Nimwegen. 2009. “Methods for Analyzing Deep Sequencing Expression Data: Constructing the Human and Mouse Promoterome with DeepCAGE Data.” Genome Biology 10 (7). https://doi.org/10.1186/gb-2009-10-7-r79.

Batada, Nizar N, and Laurence D Hurst. 2007. “Evolution of Chromosome Organization Driven by Selection for Reduced Gene Expression Noise.” Nature Genetics 39 (8): 945– 49. https://doi.org/10.1038/ng2071.

Battich, Nico, Thomas Stoeger, and Lucas Pelkmans. 2015. “Control of Transcript Variability in Single Mammalian Cells.” Cell 163: 1596–1610. https://doi.org/10.1016/j.cell.2015.11.018.

Becskei, Attila, Benjamin B Kaufmann, and Alexander van Oudenaarden. 2005. “Contributions of Low Molecule Number and Chromosomal Positioning to Stochastic Gene Expression.” Nature Genetics 37 (9): 937–44. https://doi.org/10.1038/ng1616.

Benaglia, Tatiana, Didier Chauveau, David R. Hunter, and Derek S. Young. 2009. “Mixtools: An R Package for Analyzing Finite Mixture Models.” Journal of Statistical Software 32 (6): 1–29. https://doi.org/10.18637/jss.v032.i06.

Bischl, Bernd, Michel Lang, Lars Kotthoff, Julia Schiffner, Jakob Richter, Erich Studerus, Giuseppe Casalicchio, and Zachary M. Jones. 2016. “Mlr: Machine Learning in R.” Journal of Machine Learning Research 17 (170): 1–5. http://jmlr.org/papers/v17/15-066.html.

Blake, William J, Gá Bor, Balá Zsi, Michael A Kohanski, Farren J Isaacs, Kevin F Murphy, Yina Kuang, Charles R Cantor, David R Walt, and James J Collins. 2006. “Phenotypic Consequences of Promoter-Mediated Transcriptional Noise.” Molecular Cell 24: 853–65. https://doi.org/10.1016/j.molcel.2006.11.003.

Boettiger, Alistair N, and Michael Levine. 2009. “Synchronous and Stochastic Patterns of Gene Activation in the Drosophila Embryo.” https://doi.org/10.1126/science.1173976.

Bolger, Anthony M., Marc Lohse, and Bjoern Usadel. 2014. “Trimmomatic: A Flexible Trimmer for Illumina Sequence Data.” Bioinformatics 30 (15): 2114–20. https://doi.org/10.1093/bioinformatics/btu170.

Buniello, Annalisa, Jacqueline A.L. Macarthur, Maria Cerezo, Laura W. Harris, James Hayhurst, Cinzia Malangone, Aoife McMahon, et al. 2019. “The NHGRI-EBI GWAS Catalog of Published Genome-Wide Association Studies, Targeted Arrays and Summary Statistics 2019.” Nucleic Acids Research 47 (D1): D1005–12. https://doi.org/10.1093/nar/gky1120.

Cannavò, Enrico, Nils Koelling, Dermot Harnett, David Garfield, Francesco P Casale, Lucia Ciglar, Hilary E Gustafson, et al. 2016. “Genetic Variants Regulating Expression Levels and Isoform Diversity during Embryogenesis.” Nature. https://doi.org/10.1038/nature20802.

Carey, Lucas B., David van Dijk, Peter M. A. Sloot, Jaap A. Kaandorp, and Eran Segal. 2013. “Promoter Sequence Determines the Relationship between Expression Level and Noise.” Edited by Robert Singer. PLoS Biology 11 (4): e1001528. https://doi.org/10.1371/journal.pbio.1001528.

Carninci, Piero, Albin Sandelin, Boris Lenhard, Shintaro Katayama, Kazuro Shimokawa, Jasmina Ponjavic, Colin A M Semple, et al. 2006. “Genome-Wide Analysis of Mammalian Promoter Architecture and Evolution.” Nature Genetics 38 (6): 626–35. https://doi.org/10.1038/ng1789.

Chen, Linan, Jason G Dumelie, Xiao Li, Matthew Hk Cheng, Zhiyong Yang, John D Laver, Najeeb U Siddiqui, et al. 2014. “Global Regulation of MRNA Translation and Stability in the Early Drosophila Embryo by the Smaug RNA-Binding Protein.” https://www.ncbi.nlm.nih.gov/pmc/articles/PMC4053848/pdf/gb-2014-15-1-r4.pdf.

Ciliberti, Stefano, Olivier C. Martin, Andreas Wagner, O Tenaillon, and PE Turner. 2007. “Robustness Can Evolve Gradually in Complex Regulatory Gene Networks with Varying Topology.” PLoS Computational Biology 3 (2): e15. https://doi.org/10.1371/journal.pcbi.0030015.

Crow, Megan, Nathaniel Lim, Sara Ballouz, Paul Pavlidis, and Jesse Gillis. 2019. “Predictability of Human Differential Gene Expression” 116 (13). https://doi.org/10.1073/pnas.1802973116.

Cunningham, Fiona, Premanand Achuthan, Wasiu Akanni, James Allen, M. Ridwan Amode, Irina M. Armean, Ruth Bennett, et al. 2019. “Ensembl 2019.” Nucleic Acids Research 47 (D1): D745–51. https://doi.org/10.1093/nar/gky1113.

Dempster, A. P.;, N. M.; Laird, and D.B. Rubin. 1979. “Maximum Likelihood from Incomplete Data via the EM Algorithm A.” Journal of Applied Mechanics, Transactions ASME 46 (1): 139–44. https://doi.org/10.1115/1.3424485.

Dong, Dong, Xiaojian Shao, Naiyang Deng, and Zhaolei Zhang. 2011. “Gene Expression Variations Are Predictive for Stochastic Noise.” Nucleic Acids Research 39 (2): 403–13. https://doi.org/10.1093/nar/gkq844.

Durinck, Steffen, Paul T Spellman, Ewan Birney, and Wolfgang Huber. 2009. “Mapping Identifiers for the Integration of Genomic Datasets with the R/Bioconductor Package BiomaRt.” Nature Protocols 4 (8): 1184–91. https://doi.org/10.1038/nprot.2009.97.

Edgar, Bruce A., and Patrick H. O’Farrel. 1989. “Genetic Control of Cell Division Patterns in the Drosophila Embryo.” Cell 57 (1): 177–87. https://doi.org/10.1038/jid.2014.371.

Eling, Nils, Michael D. Morgan, and John C. Marioni. 2019. “Challenges in Measuring and Understanding Biological Noise.” Nature Reviews Genetics 20 (September): 536–48. https://doi.org/10.1038/s41576-019-0130-6.

Eling, Nils, Arianne C. Richard, Sylvia Richardson, John C. Marioni, and Catalina A. Vallejos. 2018. “Correcting the Mean-Variance Dependency for Differential Variability Testing Using Single-Cell RNA Sequencing Data.” Cell Systems, August. https://doi.org/10.1016/J.CELS.2018.06.011.

Ernst, Jason, and Manolis Kellis. 2017. “Chromatin-State Discovery and Genome Annotation with ChromHMM.” Nature Protocols 12 (12): 2478–92. https://doi.org/10.1038/nprot.2017.124.

Faure, Andre J., Jörn M. Schmiedel, and Ben Lehner. 2017. “Systematic Analysis of the Determinants of Gene Expression Noise in Embryonic Stem Cells.” Cell Systems 5 (5): 471–484.e4. https://doi.org/10.1016/J.CELS.2017.10.003.

Félix, Marie-Anne, and Michalis Barkoulas. 2015. “Pervasive Robustness in Biological Systems.” Nature Publishing Group 16. https://doi.org/10.1038/nrg3949.

Filion, Guillaume J., Joke G. van Bemmel, Ulrich Braunschweig, Wendy Talhout, Jop Kind, Lucas D. Ward, Wim Brugman, et al. 2010. “Systematic Protein Location Mapping Reveals Five Principal Chromatin Types in Drosophila Cells.” Cell 143 (2): 212–24. https://doi.org/10.1016/J.CELL.2010.09.009.

Foreman, Robert, and Roy Wollman. 2019. “Mammalian Gene Expression Variability Is Explained by Underlying Cell State.” https://doi.org/10.1101/626424.

Forrest, Alistair R. R.;, Hideya; Kawaji, Michael Rehli, J. Kenneth Baillie, Michiel J. L. de Hoon, Vanja Haberle, Timo Lassmann, et al. 2014. “A Promoter-Level Mammalian Expression Atlas.” Nature 507 (7493): 462–70. https://doi.org/10.1038/nature13182.

Fraser, Hunter B, Aaron E Hirsh, Guri Giaever, Jochen Kumm, and Michael B Eisen. 2004. “Noise Minimization in Eukaryotic Gene Expression.” Edited by Ken Wolfe. PLoS Biology 2 (6): e137. https://doi.org/10.1371/journal.pbio.0020137.

Gerber, André P, Stefan Luschnig, Mark A Krasnow, Patrick O Brown, Daniel Herschlag, and Christine Guthrie. 2006. “Genome-Wide Identification of MRNAs Associated with the Translational Regulator PUMILIO in Drosophila Melanogaster.” https://www.ncbi.nlm.nih.gov/pmc/articles/PMC1400586/pdf/zpq4487.pdf.

GTEx Consortium, The GTEx. 2013. “The Genotype-Tissue Expression (GTEx) Project.” Nature Genetics 45 (6): 580–85. https://doi.org/10.1038/ng.2653.

Haberle, Vanja, Alistair R.R. Forrest, Yoshihide Hayashizaki, Piero Carninci, and Boris Lenhard. 2015. “CAGEr: Precise TSS Data Retrieval and High-Resolution Promoterome Mining for Integrative Analyses.” Nucleic Acids Research 43 (8). https://doi.org/10.1093/nar/gkv054.

Haberle, Vanja, and Alexander Stark. 2018. “Eukaryotic Core Promoters and the Functional Basis of Transcription Initiation.” Nature Reviews Molecular Cell Biology, June, 1. https://doi.org/10.1038/s41580-018-0028-8.

Haeussler, Maximilian, Ann S. Zweig, Cath Tyner, Matthew L. Speir, Kate R. Rosenbloom, Brian J. Raney, Christopher M. Lee, et al. 2019. “The UCSC Genome Browser Database: 2019 Update.” Nucleic Acids Research 47 (D1): D853–58. https://doi.org/10.1093/nar/gky1095.

Hammonds, Ann S, Christopher A Bristow, William W Fisher, Richard Weiszmann, Siqi Wu, Volker Hartenstein, Manolis Kellis, Bin Yu, Erwin Frise, and Susan E Celniker. 2013. “Spatial Expression of Transcription Factors in Drosophila Embryonic Organ Development.” https://www.ncbi.nlm.nih.gov/pmc/articles/PMC4053779/pdf/gb-2013-14-12-r140.pdf.

Hart, Traver, Amy Hin Yan Tong, Katie Chan, Jolanda Van Leeuwen, Ashwin Seetharaman, Michael Aregger, Megha Chandrashekhar, et al. 2017. “Evaluation and Design of Genome-Wide CRISPR/SpCas9 Knockout Screens.” G3: Genes, Genomes, Genetics 7 (8): 2719–27. https://doi.org/10.1534/g3.117.041277.

Hoskins, Roger A, Jane M Landolin, James B Brown, Jeremy E Sandler, Hazuki Takahashi, Timo Lassmann, Charles Yu, et al. 2011. “Genome-Wide Analysis of Promoter Architecture in Drosophila Melanogaster.” Genome Research 21 (2): 182–92. https://doi.org/10.1101/gr.112466.110.

Hu, Yanhui, Ian Flockhart, Arunachalam Vinayagam, Clemens Bergwitz, Bonnie Berger, Norbert Perrimon, and Stephanie E. Mohr. 2011. “An Integrative Approach to Ortholog Prediction for Disease-Focused and Other Functional Studies.” BMC Bioinformatics 12. https://doi.org/10.1186/1471-2105-12-357.

Huang, Sui. 2009. “Non-Genetic Heterogeneity of Cells in Development: More than Just Noise.” Development (Cambridge, England) 136 (23): 3853–62. https://doi.org/10.1242/dev.035139.

Huang, Wen, Andreas Massouras, Yutaka Inoue, Jason Peiffer, Miquel Ràmia, Aaron M. Tarone, Lavanya Turlapati, et al. 2014. “Natural Variation in Genome Architecture among 205 Drosophila Melanogaster Genetic Reference Panel Lines.” Genome Research 24 (7): 1193–1208. https://doi.org/10.1101/gr.171546.113.

Junion, Guillaume, Mikhail Spivakov, Charles Girardot, Martina Braun, E Hilary Gustafson, Ewan Birney, and Eileen E M Furlong. 2012. “A Transcription Factor Collective Defines Cardiac Cell Fate and Reflects Lineage History.” https://doi.org/10.1016/j.cell.2012.01.030.

Kaneko, Kunihiko. 2011. “Proportionality between Variances in Gene Expression Induced by Noise and Mutation: Consequence of Evolutionary Robustness.” BMC Evolutionary Biology 11 (1): 27. https://doi.org/10.1186/1471-2148-11-27.

Kedlian, Veronika R, Handan Melike Donertas, and Janet M Thornton. 2019. “The Variability of Expression of Many Genes and Most Functional Pathways Is Observed to Increase with Age in Brain Transcriptome Data.” https://doi.org/10.1101/526491.

Kent, W. J., C. W. Sugnet, T. S. Furey, K. M. Roskin, T. H. Pringle, A. M. Zahler, and a. D. Haussler. 2002. “The Human Genome Browser at UCSC.” Genome Research 12 (6): 996–1006. https://doi.org/10.1101/gr.229102.

Khan, Aziz, Oriol Fornes, Arnaud Stigliani, Marius Gheorghe, Jaime A. Castro-Mondragon, Robin Van Der Lee, Adrien Bessy, et al. 2018. “JASPAR 2018: Update of the Open-Access Database of Transcription Factor Binding Profiles and Its Web Framework.” Nucleic Acids Research 46 (D1): D260–66. https://doi.org/10.1093/nar/gkx1126.

Kozomara, Ana, Maria Birgaoanu, and Sam Griffiths-Jones. 2019. “MiRBase: From MicroRNA Sequences to Function.” Nucleic Acids Research 47 (D1): D155–62. https://doi.org/10.1093/nar/gky1141.

Kozomara, Ana, and Sam Griffiths-Jones. 2014. “MiRBase: Annotating High Confidence MicroRNAs Using Deep Sequencing Data.” Nucleic Acids Research 42 (D1): 68–73. https://doi.org/10.1093/nar/gkt1181.

Kudron, Michelle M., Alec Victorsen, Louis Gevirtzman, LaDeana W. Hillier, William W. Fisher, Dionne Vafeados, Matt Kirkey, et al. 2018. “The ModERN Resource: Genome-Wide Binding Profiles for Hundreds of *Drosophila* and *Caenorhabditis Elegans* Transcription Factors.” Genetics 208 (3): 937–49. https://doi.org/10.1534/genetics.117.300657.

Kursa, Miron B, and Witold R Rudnicki. 2010. “Feature Selection with the Boruta Package.” JSS Journal of Statistical Software. Vol. 36. http://www.jstatsoft.org/.

Kvon, Evgeny Z, Tomas Kazmar, Gerald Stampfel, J Omar Yáñez-Cuna, Michaela Pagani, Katharina Schernhuber, Barry J Dickson, and Alexander Stark. 2014. “Genome-Scale Functional Characterization of Drosophila Developmental Enhancers in Vivo.” Nature 512. https://doi.org/10.1038/nature13395.

Larsson, Anton J. M., Per Johnsson, Michael Hagemann-Jensen, Leonard Hartmanis, Omid R. Faridani, Björn Reinius, Åsa Segerstolpe, Chloe M. Rivera, Bing Ren, and Rickard Sandberg. 2019. “Genomic Encoding of Transcriptional Burst Kinetics.” Nature 565 (7738): 251–54. https://doi.org/10.1038/s41586-018-0836-1.

Lawrence, Michael, Wolfgang Huber, Hervé Pagès, Patrick Aboyoun, Marc Carlson, Robert Gentleman, Martin T. Morgan, and Vincent J. Carey. 2013. “Software for Computing and Annotating Genomic Ranges.” Edited by Andreas Prlic. PLoS Computational Biology 9 (8): e1003118. https://doi.org/10.1371/journal.pcbi.1003118.

Lehner, Ben. 2008. “Selection to Minimise Noise in Living Systems and Its Implications for the Evolution of Gene Expression.” Molecular Systems Biology 4: 170. https://doi.org/10.1038/msb.2008.11.

Lenhard, Boris, Albin Sandelin, and Piero Carninci. 2012. “Metazoan Promoters: Emerging Characteristics and Insights into Transcriptional Regulation.” Nature Reviews Genetics 13 (4): 233–45. https://doi.org/10.1038/nrg3163.

Li, H., B. Handsaker, A. Wysoker, T. Fennell, J. Ruan, N. Homer, G. Marth, G. Abecasis, and R. Durbin. 2009. “The Sequence Alignment/Map Format and SAMtools.” Bioinformatics 25 (16): 2078–79. https://doi.org/10.1093/bioinformatics/btp352.

Li, Heng, and Richard Durbin. 2010. “Fast and Accurate Long-Read Alignment with Burrows-Wheeler Transform.” Bioinformatics 26 (5): 589–95. https://doi.org/10.1093/bioinformatics/btp698.

Lippert, Christoph, Francesco Paolo Casale, Barbara Rakitsch, and Oliver Stegle. 2014. “LIMIX: Genetic Analysis of Multiple Traits.” BioRxiv, May, 003905. https://doi.org/10.1101/003905.

Liu, Jialin, Michael Frochaux, Vincent Gardeux, Bart Deplancke, and Marc Robinson- Rechavi. 2019. “Selection against Expression Noise Explains the Origin of the Hourglass Pattern of Evo-Devo.” BioRxiv, 700997. https://doi.org/10.1101/700997.

Lizio, Marina, Jayson Harshbarger, Hisashi Shimoji, Jessica Severin, Takeya Kasukawa, Serkan Sahin, Imad Abugessaisa, et al. 2015. “Gateways to the FANTOM5 Promoter Level Mammalian Expression Atlas.” Genome Biology 16 (1): 1–14. https://doi.org/10.1186/s13059-014-0560-6.

Love, M. I., Simon Anders, and Wolfgang Huber. 2014. Differential Analysis of Count Data - the DESeq2 Package. Genome Biology. Vol. 15. https://doi.org/110.1186/s13059-014-0550-8.

Mackay, Trudy F. C., Stephen Richards, Eric A. Stone, Antonio Barbadilla, Julien F. Ayroles, Dianhui Zhu, Sònia Casillas, et al. 2012. “The Drosophila Melanogaster Genetic Reference Panel.” Nature 482 (7384): 173–78. https://doi.org/10.1038/nature10811.

Macneil, Lesley T, and Albertha J M Walhout. 2011. “Gene Regulatory Networks and the Role of Robustness and Stochasticity in the Control of Gene Expression.” https://doi.org/10.1101/gr.097378.109.

Metzger, Brian P. H., David C. Yuan, Jonathan D. Gruber, Fabien Duveau, and Patricia J. Wittkopp. 2015. “Selection on Noise Constrains Variation in a Eukaryotic Promoter.” Nature 521 (7552): 344–47. https://doi.org/10.1038/nature14244.

Morgan, Michael D., and John C. Marioni. 2018. “CpG Island Composition Differences Are a Source of Gene Expression Noise Indicative of Promoter Responsiveness.” Genome Biology 19 (1): 81. https://doi.org/10.1186/s13059-018-1461-x.

Moskalev, Alexey, Svetlana Zhikrivetskaya, George Krasnov, Mikhail Shaposhnikov, Ekaterina Proshkina, Dmitry Borisoglebsky, Anton Danilov, et al. 2015. “A Comparison of the Transcriptome of Drosophila Melanogaster in Response to Entomopathogenic Fungus, Ionizing Radiation, Starvation and Cold Shock.” BMC Genomics 16 Suppl 1 (Suppl 13): S8. https://doi.org/10.1186/1471-2164-16-S13-S8.

Nelson, Matthew R, Daniel Wegmann, Margaret G Ehm, Darren Kessner, Pamela St, Claudio Verzilli, Judong Shen, et al. 2015. “An Abundance of Rare Functional Variants in 202 Drug Target Genes Sequenced in 14,002 People.” Science 337 (6090): 100–104. https://doi.org/10.1126/science.1217876.An.

Ohler, Uwe. 2006. “Identification of Core Promoter Modules in Drosophila and Their Application in Accurate Transcription Start Site Prediction.” Nucleic Acids Research 34 (20): 5943–50. https://doi.org/10.1093/nar/gkl608.

Ohler, Uwe, Guo-chun Liao, Heinrich Niemann, and Gerald M Rubin. 2002. “Computational Analysis of Core Promoters in the Drosophila Genome.” Genome Biology 3 (12): RESEARCH0087. https://doi.org/10.1186/GB-2002-3-12-RESEARCH0087.

Perry, Michael W, Alistair N Boettiger, Jacques P Bothma, and Michael Levine. 2010. “Shadow Enhancers Foster Robustness of Drosophila Gastrulation.” Current Biology : CB 20 (17): 1562–67. https://doi.org/10.1016/j.cub.2010.07.043.

Quinlan, Aaron R., and Ira M. Hall. 2010. “BEDTools: A Flexible Suite of Utilities for Comparing Genomic Features.” Bioinformatics 26 (6): 841–42. https://doi.org/10.1093/bioinformatics/btq033.

R Development Core Team. 2013. “A Language and Environment for Statistical Computing.” R Foundation for Statistical Computing. http://www.r-project.org.

Rach, Elizabeth A, Hsiang-Yu Yuan, William H Majoros, Pavel Tomancak, and Uwe Ohler. 2009. “Motif Composition, Conservation and Condition-Specificity of Single and Alternative Transcription Start Sites in the Drosophila Genome.” Genome Biology 10 (7). https://doi.org/10.1186/gb-2009-10-7-r73.

Raj, Arjun, and Alexander van Oudenaarden. 2008. “Nature, Nurture, or Chance: Stochastic Gene Expression and Its Consequences.” Cell 135 (2): 216–26. https://doi.org/10.1016/j.cell.2008.09.050.

Ramírez, Fidel, Vivek Bhardwaj, Laura Arrigoni, Kin Chung Lam, Björn A Grüning, José Villaveces, Bianca Habermann, Asifa Akhtar, and Thomas Manke. 2018. “High-Resolution TADs Reveal DNA Sequences Underlying Genome Organization in Flies.” https://doi.org/10.1038/s41467-017-02525-w.

Ran, Di, and Z John Daye. 2017. “Gene Expression Variability and the Analysis of Large-Scale RNA-Seq Studies with the MDSeq.” Nucleic Acids Research 45 (13): e127. https://doi.org/10.1093/nar/gkx456.

Raser, J M, and E K O’Shea. 2005. “Noise in Gene Expression: Orgins, Consequences, and Control.” Science 309 (5743): 2010–13. https://doi.org/10.1126/science.1105891.

Ravarani, Charles N J, Guilhem Chalancon, Michal Breker, Natalia Sanchez De Groot, and M Madan Babu. 2015. “Affinity and Competition for TBP Are Molecular Determinants of Gene Expression Noise.” https://doi.org/10.1038/ncomms10417.

Ray, Debashish, Hilal Kazan, Kate B. Cook, Matthew T. Weirauch, Hamed S. Najafabadi, Xiao Li, Serge Gueroussov, et al. 2013. “A Compendium of RNA-Binding Motifs for Decoding Gene Regulation.” Nature 499 (7457): 172–77. https://doi.org/10.1038/nature12311.

Richard, Angélique, Loïs Boullu, Ulysse Herbach, Arnaud Bonnafoux, Valérie Morin, Elodie Vallin, Anissa Guillemin, et al. 2016. “Single-Cell-Based Analysis Highlights a Surge in Cell-to-Cell Molecular Variability Preceding Irreversible Commitment in a Differentiation Process.” Edited by Sarah A. Teichmann. PLOS Biology 14 (12): e1002585. https://doi.org/10.1371/journal.pbio.1002585.

Saunders, Abbie, Leighton J Core, Catherine Sutcliffe, John T Lis, and Hilary L Ashe. 2013. “Extensive Polymerase Pausing during Drosophila Axis Patterning Enables High-Level and Pliable Transcription.” https://doi.org/10.1101/gad.215459.113.

Schmiedel, Jörn M, Sandy L Klemm, Yannan Zheng, Apratim Sahay, Nils Blüthgen, Debora S Marks, and Alexander van Oudenaarden. 2015. “Gene Expression. MicroRNA Control of Protein Expression Noise.” Science (New York, N.Y.) 348 (6230): 128–32. https://doi.org/10.1126/science.aaa1738.

Schmiedel, Jörn M, Debora S Marks, Ben Lehner, and Nils Blüthgen. 2018. “Noise Control Is a Primary Function of MicroRNAs and Post-Transcriptional Regulation.” https://doi.org/10.1101/168641.

Schor, Ignacio E, Jacob F Degner, Dermot Harnett, Enrico Cannavò, Francesco P Casale, Heejung Shim, David A Garfield, et al. 2017. “Promoter Shape Varies across Populations and Affects Promoter Evolution and Expression Noise.” Nature Publishing Group 49. https://doi.org/10.1038/ng.3791.

Tomancak, Pavel, Amy Beaton, Richard Weiszmann, Elaine Kwan, Sheng Qiang Shu, Suzanna E. Lewis, Stephen Richards, et al. 2002. “Systematic Determination of Patterns of Gene Expression during Drosophila Embryogenesis.” Genome Biology 3 (12): 1–14. https://doi.org/10.1186/gb-2002-3-12-research0088.

Viñuela, Ana, Andrew A Brown, Alfonso Buil, Pei-Chien Tsai, Matthew N Davies, Jordana T Bell, Emmanouil T Dermitzakis, Timothy D Spector, and Kerrin S Small. 2018. “Age-Dependent Changes in Mean and Variance of Gene Expression across Tissues in a Twin Cohort.” Human Molecular Genetics 27 (4): 732–41. https://doi.org/10.1093/hmg/ddx424.

Vorontsov, Ilya E., Alla D. Fedorova, Ivan S. Yevshin, Ruslan N. Sharipov, Fedor A. Kolpakov, Vsevolod J. Makeev, and Ivan V. Kulakovskiy. 2018. “Genome-Wide Map of Human and Mouse Transcription Factor Binding Sites Aggregated from ChIP-Seq Data.” BMC Research Notes 11 (1): 10–12. https://doi.org/10.1186/s13104-018-3856-x.

Waddington, CH H. 1942. “Canalization of Development and the Inheritance of Acquired Characters.” Nature 150 (3811): 563–65. https://doi.org/10.1038/150563a0.

Weirauch, Matthew T, Ally Yang, Mihai Albu, Atina Cote, Alejandro Montenegro-, Philipp Drewe, Hamed S Najafabadi, et al. 2015. “Determination and Inference of Eukaryotic Transcription Factor Sequence Specificity” 158 (6): 1431–43. https://doi.org/10.1016/j.cell.2014.08.009.Determination.

Wishart, David S., Yannick D. Feunang, An C. Guo, Elvis J. Lo, Ana Marcu, Jason R. Grant, Tanvir Sajed, et al. 2018. “DrugBank 5.0: A Major Update to the DrugBank Database for 2018.” Nucleic Acids Research 46 (D1): D1074–82. https://doi.org/10.1093/nar/gkx1037.

Wright, Marvin N, and Andreas Ziegler. 2015. “Ranger: A Fast Implementation of Random Forests for High Dimensional Data in C++ and R.” https://arxiv.org/pdf/1508.04409.pdf.

Yu, Guangchuang, Li Gen Wang, Yanyan Han, and Qing Yu He. 2012. “ClusterProfiler: An R Package for Comparing Biological Themes among Gene Clusters.” OMICS A Journal of Integrative Biology 16 (5): 284–87. https://doi.org/10.1089/omi.2011.0118.

Zhang, Fuquan, Yin Yao Shugart, Weihua Yue, Zaohuo Cheng, Guoqiang Wang, Zhenhe Zhou, Chunhui Jin, Jianmin Yuan, Sha Liu, and Yong Xu. 2015. “Increased Variability of Genomic Transcription in Schizophrenia.” Scientific Reports 5 (December): 17995. https://doi.org/10.1038/srep17995.

Zinzen, Robert P, Charles Girardot, Julien Gagneur, Martina Braun, and Eileen E M Furlong. 2009. “Combinatorial Binding Predicts Spatio-Temporal Cis-Regulatory Activity.” Nature 461. https://doi.org/10.1038/nature08531.

