## Supplementary Figures and Supplementary Tables legends for "Predictive features of gene expression variation reveal a mechanistic link between expression variation and differential expression"

**Supplementary Material:**

- **Supplementary Figures 1-8**
- **Supplementary Tables 1-17**

Fig. S1

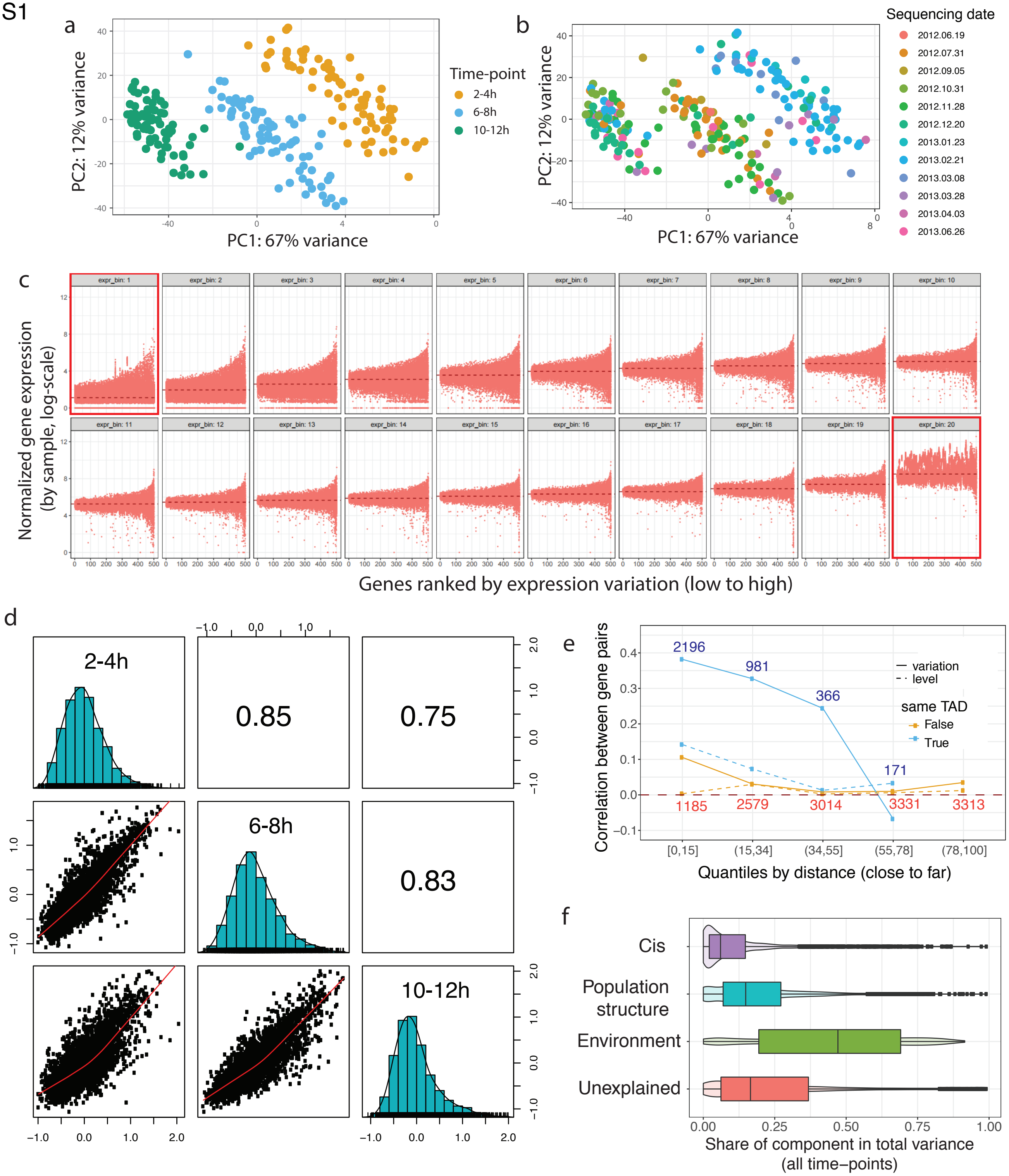

a
PC2: 12% variance
PC1: 67% variance
Time-point
2-4h
6-8h
10-12h
b
Sequencing date
2012.06.19
2012.07.31
2012.09.05
2012.10.31
2012.11.28
2012.12.20
2013.01.23
2013.02.21
2013.03.08
2013.03.28
2013.04.03
2013.06.26
c
Normalized gene expression
(by sample, log-scale)
Genes ranked by expression variation (low to high)
expr_bin: 1
expr_bin: 20
d
2-4h
6-8h
10-12h
0.85
0.75
0.83
e
Correlation between gene pairs
Quantiles by distance (close to far)
variation
level
same TAD
False
True
2196
981
366
171
1185
2579
3014
3331
3313
[0,15]
(15,34]
(34,55]
(55,78]
(78,100]
f
Cis
Population structure
Environment
Unexplained
Share of component in total variance
(all time–points)

**Supplementary Figure 1. (a-b**) Samples cluster by developmental time-points (a) and not by sequencing batches (b). First two principal components for 225 3' Tag-seq samples (75 samples at three time-points). Each dot represents a sample. Samples are colored by time-points (a) and sequencing dates. All genes with quantified gene expression were used for the analysis. PCA was done on the raw expression counts after applying variance stabilization transformation from DESeq2 (Anders and Huber 2010). **(c)** Residual expression variation reflects expression heterogeneity (across samples) at any given expression level. Gene expression across samples (y-xis, size factor normalized read counts) for genes binned by median expression level (20 subplots for 5%-percentiles by expression level) and ordered by expression variation (x-axis, residual coefficient of variation). Each dot represents a gene in one of 75 samples. Top and bottom 5% of genes by expression level were removed as potential source of outliers (the first and the last sub-plots, outlined with red).Data is shown for 10-12h time-point. **(d)** Gene expression variation is consistent across timepoints. Diagonal: distributions of gene expression variation values at three time-points. Upper triangular panels: Spearman correlation coefficients of gene expression variation between pairs of time-points. Lower triangular panels: scatter plots showing correlation between gene expression variation at different time-points (x and y-axis). Each dot represents a gene. Final set of 4074 genes was used at all three time-points. **(e)** Correlation in expression variation (solid lines) and median expression levels (dashed lines) between genes located at varying quantile distances between their TSSs (actual distance intervals in kB are shown on the x-axis). Only gene pairs located on the same chromosome and with TSS-to-TSS distance <100 kB are considered. Spearman correlation coefficient (y-axis) and quantiles by distances (in kB) (x-axis) is plotted separately for gene pairs within the same TAD (blue) or split into different TADs (orange). Number of gene pairs in each group is indicated (blue and orange font for gene pairs within the same TAD or crossing TAD border, respectively), only groups with >100 gene pairs are shown. **(f)** Share of total variance (across 225 samples) of genes explained by different components according to LIMIX variance decomposition (Methods). All three time-points were used in variance decomposition to achieve better convergence of the algorithm and more precise estimate of Cis component. Results for the final set of 4074 genes are plotted. Only Cis component (defined here as sum of Cis and (Cis x Environment) components from LIMIX) was used as a feature in subsequent random forest models.

Fig. S2

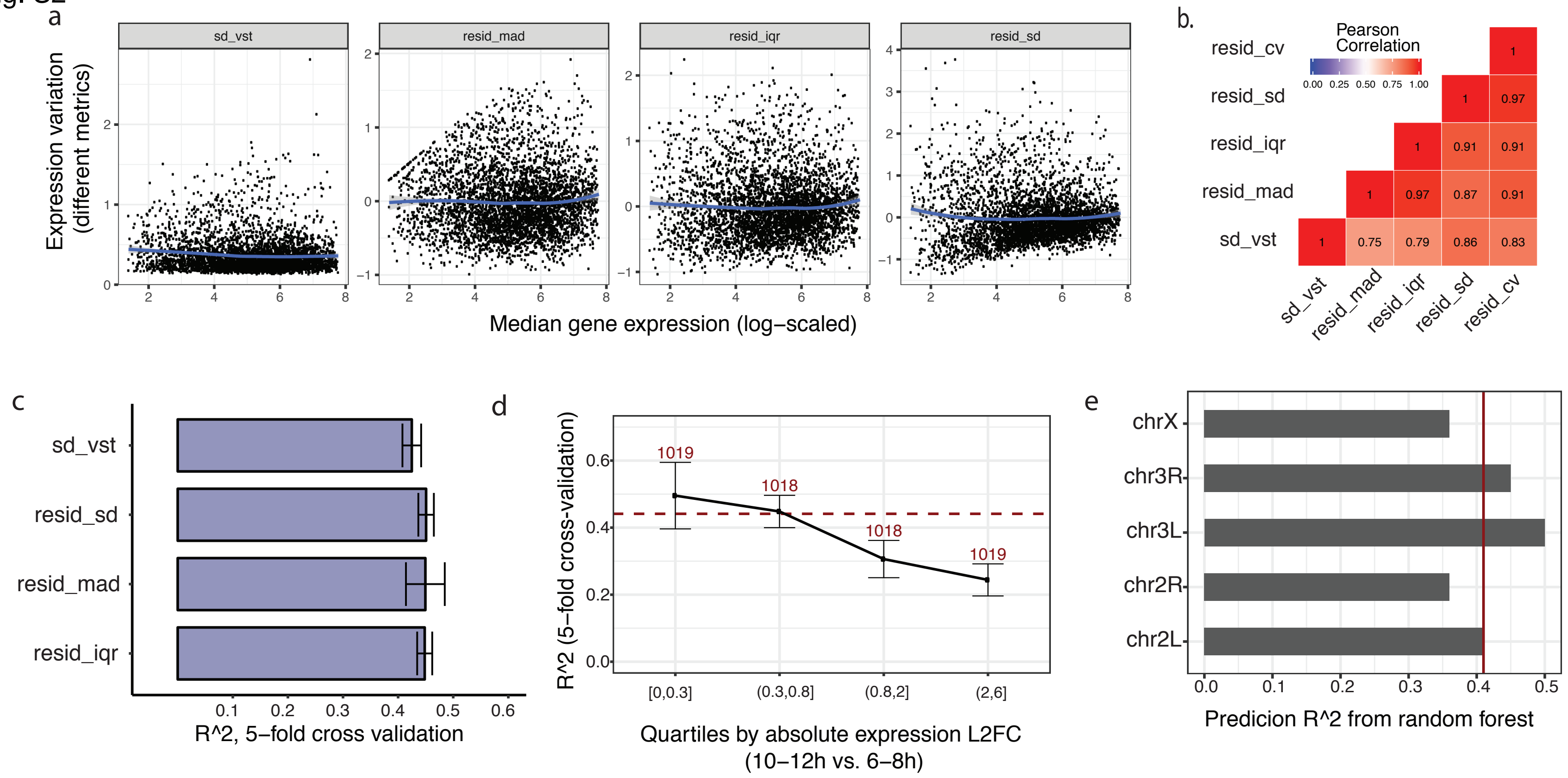

a
sd_vst
resid_mad
resid_iqr
resid_sd
Expression variation (different metrics)
Median gene expression (log−scaled)
b.
Pearson Correlation
0.00 0.25 0.50 0.75 1.00
resid_cv
resid_sd
resid_iqr
resid_mad
sd_vst
1
1 0.97
1 0.91 0.91
1 0.97 0.87 0.91
1 0.75 0.79 0.86 0.83
sd_vst resid_mad resid_iqr resid_sd resid_cv
c
sd_vst
resid_sd
resid_mad
resid_iqr
0.1 0.2 0.3 0.4 0.5 0.6
R^2, 5−fold cross validation
d
R^2 (5−fold cross-validation)
1019
1018
1018
1019
[0,0.3]
(0.3,0.8]
(0.8,2]
(2,6]
Quartiles by absolute expression L2FC
(10−12h vs. 6−8h)
e
chrX
chr3R
chr3L
chr2R
chr2L
0.0 0.1 0.2 0.3 0.4 0.5
Predicion R^2 from random forest

**Supplementary Figure 2. (a)** Alternative measures of gene expression variation corrected for level dependence. From left to right: standard deviation after variance stabilizing transformation (*sd_vst*) from DEseq2 (Anders and Huber 2010); residual median absolute deviation (resid_mad), inter-quartile range (resid_iqr), and residual standard deviation (resid_sd). The three later measures are LOESS residuals from regression on median expression level calculated in the same way as residual coefficient of variation (Methods). Data is shown for the final set of 4074 genes at 10-12h. Each dot represents a gene. Blue line shows LOESS regression fit, indicating no global dependence between corrected variation measures and median gene expression level. **(b)** Spearman correlation coefficient among different expression variation measures. Final variation measure used in the analysis is residual coefficient of variation (resid_cv). Correlations were computed on the final set of 4074 genes at 10-12h. **(c)** Random forest performance (R^2 from 5-fold cross-validation) for alternative measures of expression variation (same as in a.). Random forest was run on the set of features important for predicting expression variation (residual coefficient of variation, same as in Fig 1d). Whiskers correspond to standard deviation across the 5-fold cross validation. **(d)** Performance of random forest (R^2 from 5-fold cross-validation) for predicting expression variation for genes split into four quartiles by expression log2-fold change between 10-12h and 6-8h time-points. Numbers on the plot indicate number of genes in the corresponding quartiles. Horizontal red line indicates model performance on the full dataset (as in Fig 1d). Whiskers correspond to standard deviation across the 5-fold cross validation. **(e)** Performance of random forest for predicting expression variation when test and train sets come from different chromosome (arms). Prediction R^2 for each chromosome (arm) are shown as horizontal bars. Vertical red line indicates median performance across chromosomes. Whiskers correspond to standard deviation across the 5-fold cross validation.

Fig. S3

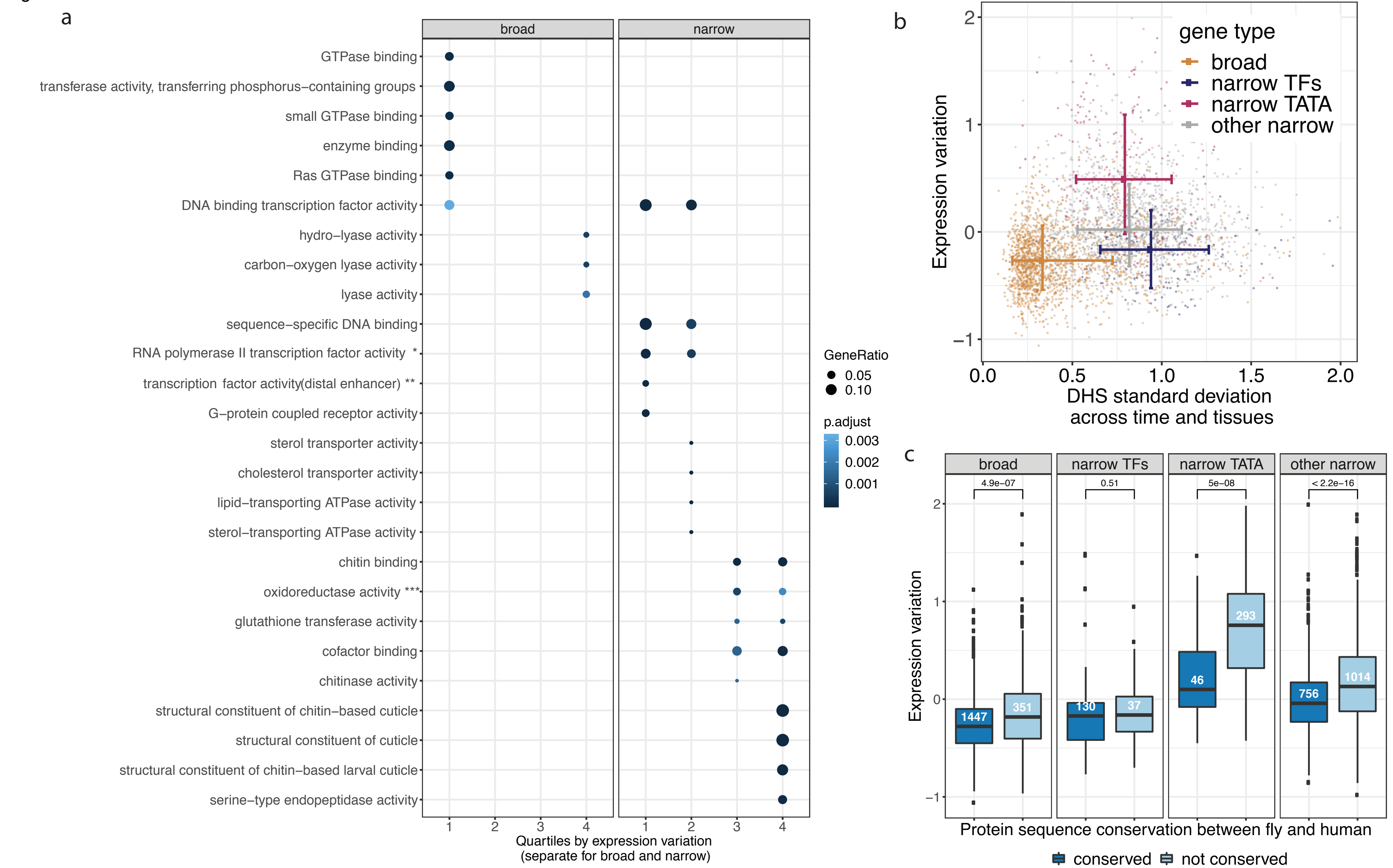
a
broad
narrow
GTPase binding
transferase activity, transferring phosphorus−containing groups
small GTPase binding
enzyme binding
Ras GTPase binding
DNA binding transcription factor activity
hydro−lyase activity
carbon−oxygen lyase activity
lyase activity
sequence−specific DNA binding
RNA polymerase II transcription factor activity *
transcription factor activity(distal enhancer) **
G−protein coupled receptor activity
sterol transporter activity
cholesterol transporter activity
lipid−transporting ATPase activity
sterol−transporting ATPase activity
chitin binding
oxidoreductase activity ***
glutathione transferase activity
cofactor binding
chitinase activity
structural constituent of chitin−based cuticle
structural constituent of cuticle
structural constituent of chitin−based larval cuticle
serine−type endopeptidase activity
Quartiles by expression variation
(separate for broad and narrow)
GeneRatio
0.05
0.10
p.adjust
0.003
0.002
0.001
b
gene type
broad
narrow TFs
narrow TATA
other narrow
Expression variation
DHS standard deviation
across time and tissues
c
broad
narrow TFs
narrow TATA
other narrow
4.9e−07
0.51
5e−08
< 2.2e−16
1447
351
130
37
46
293
756
1014
Expression variation
Protein sequence conservation between fly and human
conserved
not conserved

**Supplementary Figure 3. (a)** GO functional enrichment (Molecular function) of genes with broad (left panel) and narrow promoters (right panel) split into four quantiles by expression variation (x-axis). Quartiles of expression variation (1- lowest, 4 – highest, same as in Fig. 3c) were calculated for broad and narrow promoter genes separately. Quantile intervals for broad promoter genes (1 to 4): [-1.06,-0.444]; (-0.444,-0.266]; (-0.266,-0.0754]; (-0.0754,1.89]. Quantile intervals for narrow promoter genes (1 to 4): [-0.98,-0.173]; (-0.173,0.0751]; (0.0751,0.416]; (0.416,1.99]. **(b)** Developmental variation (standard deviation across tissues and developmental time-points) of DHSs (x-axis) around TSSs of genes with different expression variation (y-axis). Each DHS was assigned to the closest gene for this analysis, only DHSs located less than 500 bp. from the annotated TSSs are considered. Median of corresponding DHS standard deviations was calculated for each gene (x-axis) and plotted against gene expression variation. Each dot represents a gene. Colors represent types of the corresponding genes (same as in Fig. 3a). Whiskers indicate one standard deviation around mean value for the corresponding groups of genes. **(c)** Expression variation for different groups of genes (same as in b.) dependent on whether they have orthologs in human. Only orthologs with high conservation rank were considered (Methods). Wilcoxon test p-values are shown above the boxplots.

Fig. S4

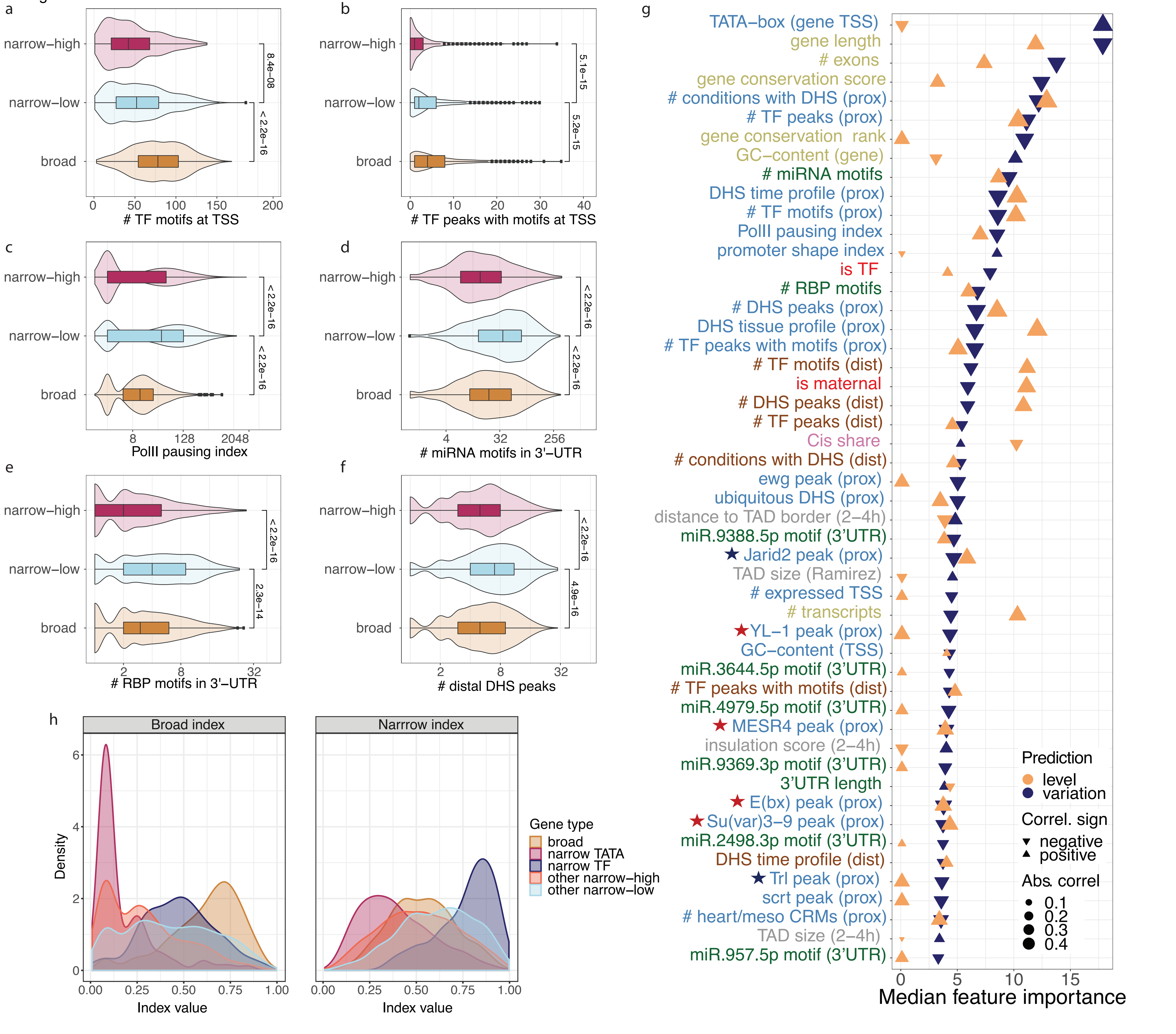
a
narrow–high
narrow–low
broad
8.4e–08
< 2.2e–16
0
50
100
150
200
### TF motifs at TSS
b
narrow–high
narrow–low
broad
5.1e–15
5.2e–15
0
10
20
30
40
### TF peaks with motifs at TSS
c
narrow–high
narrow–low
broad
< 2.2e–16
< 2.2e–16
8
128
2048
PolII pausing index
d
narrow–high
narrow–low
broad
< 2.2e–16
< 2.2e–16
4
32
256
### miRNA motifs in 3'–UTR
e
narrow–high
narrow–low
broad
< 2.2e–16
2.3e–14
2
8
32
### RBP motifs in 3'–UTR
f
narrow–high
narrow–low
broad
< 2.2e–16
4.9e–16
2
8
32
### distal DHS peaks
h
Broad index
Narrrow index
Density
0
2
4
6
0.00
0.25
0.50
0.75
1.00
Index value
Gene type
broad
narrow TATA
narrow TF
other narrow–high
other narrow–low
g
TATA–box (gene TSS)
gene length
### exons
gene conservation score
### conditions with DHS (prox)
### TF peaks (prox)
gene conservation rank
GC–content (gene)
### miRNA motifs
DHS time profile (prox)
### TF motifs (prox)
PolII pausing index
promoter shape index
is TF
### RBP motifs
### DHS peaks (prox)
DHS tissue profile (prox)
### TF peaks with motifs (prox)
### TF motifs (dist)
is maternal
### DHS peaks (dist)
### TF peaks (dist)
Cis share
### conditions with DHS (dist)
ewg peak (prox)
ubiquitous DHS (prox)
distance to TAD border (2–4h)
miR.9388.5p motif (3'UTR)
★ Jarid2 peak (prox)
TAD size (Ramirez)
### expressed TSS
### transcripts
★ YL–1 peak (prox)
GC–content (TSS)
miR.3644.5p motif (3'UTR)
### TF peaks with motifs (dist)
miR.4979.5p motif (3'UTR)
★ MESR4 peak (prox)
insulation score (2–4h)
miR.9369.3p motif (3'UTR)
3'UTR length
★ E(bx) peak (prox)
★ Su(var)3–9 peak (prox)
miR.2498.3p motif (3'UTR)
DHS time profile (dist)
★ Trl peak (prox)
scrt peak (prox)
### heart/meso CRMs (prox)
TAD size (2–4h)
miR.957.5p motif (3'UTR)
0
5
10
15
Median feature importance
Prediction
level
variation
Correl. sign
negative
positive
Abs. correl
0.1
0.2
0.3
0.4

**Supplementary Figure 4**

**(a-f)**. Differences between broad, narrow-low and narrow-high genes by the number of different TF motifs in TSS-proximal DHSs **(a)**, number of different TF peaks with motifs in TSS-proximal DHSs **(b),** polymerase II pausing index **(c)**, number of different miRNA motifs in 3'UTR **(d)**, number of different RNA-binding protein motifs in 3'UTRs **(e)**, and number of TSS-distal DHSs (more than 500 bp and less than 10 kB around TSS) **(f)**. P-values come from Wilcoxon rank test. **(g)** Top-50 important features for predicting expression variation within narrow promoter genes according to Boruta feature selection algorithm (with the corresponding importance for predicting expression level). Features are ordered by their importance for expression variation. Blue triangles indicate importance for variation, orange for level. Size and orientation of triangles correspond to absolute value and sign of correlation coefficient of feature with predicted variable, respectively. For binary features, point-biserial coefficient of correlation was used, otherwise Spearman coefficient of correlation. Label colors correspond to feature groups (same as in Fig. 2f).

Fig. S5

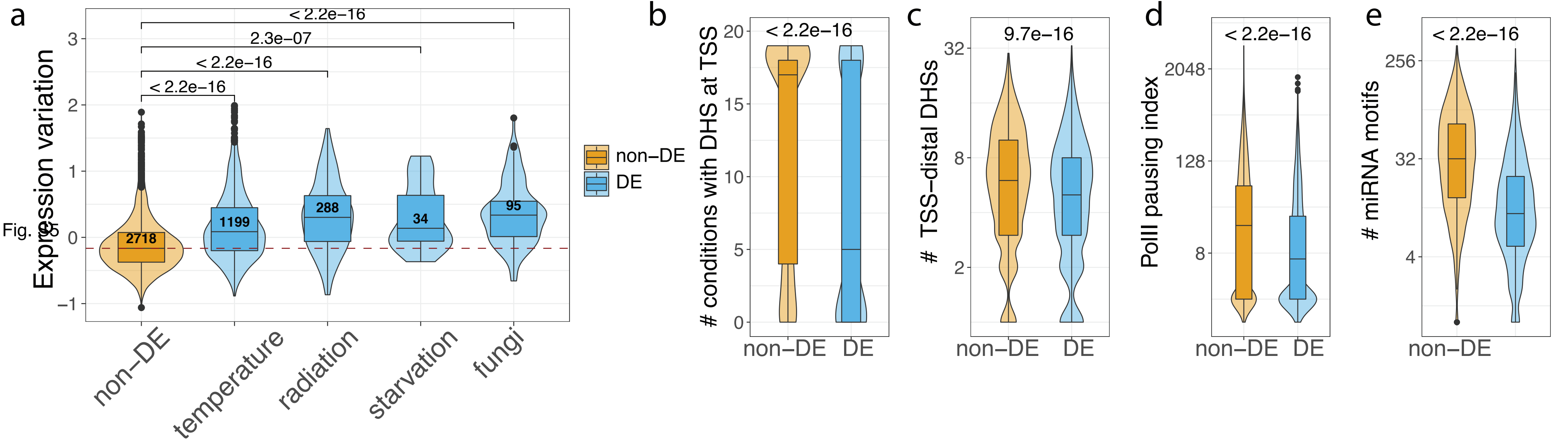

a
Expression variation
< 2.2e−16
2.3e−07
< 2.2e−16
< 2.2e−16
2718
1199
288
34
95
non−DE
temperature
radiation
starvation
fungi
non−DE
DE
b
### conditions with DHS at TSS
< 2.2e−16
non−DE
DE
c
### TSS−distal DHSs
9.7e−16
non−DE
DE
d
PolII pausing index
< 2.2e−16
non−DE
DE
e
### miRNA motifs
< 2.2e−16
non−DE

**Supplementary Figure 5**

**(a)** Expression variation (in our embryonic dataset) of genes differentially expressed (DE) upon different stress conditions from (Moskalev et al. 2015) compared to genes not differentially expressed in any of these experiments (non-DE). Stress conditions include: temperature (union of DE genes in three cold shock experiments at +4$^{o}$C, 0$^{o}$C, -4$^{o}$C), radiation (union of 144 Gly, 360 Gly, and 864 Gly ionizing radiation), starvation (16 h), and fungi (union of 10 CFU and 100 CFU entomopathogenic fungus infection). P-values come from Wilcoxon rank test. **(b-e)** Differences between DE genes (genes differentially expression in at least one stress condition from above) and non-DE genes by the number of conditions with DHS at TSS **(b),** and number of TSS-distal DHSs **(c),** polymerase II pausing index **(d)**, number of different miRNA motifs in 3’ UTR **(e)**. P-values come from Wilcoxon rank test.

Fig. S6

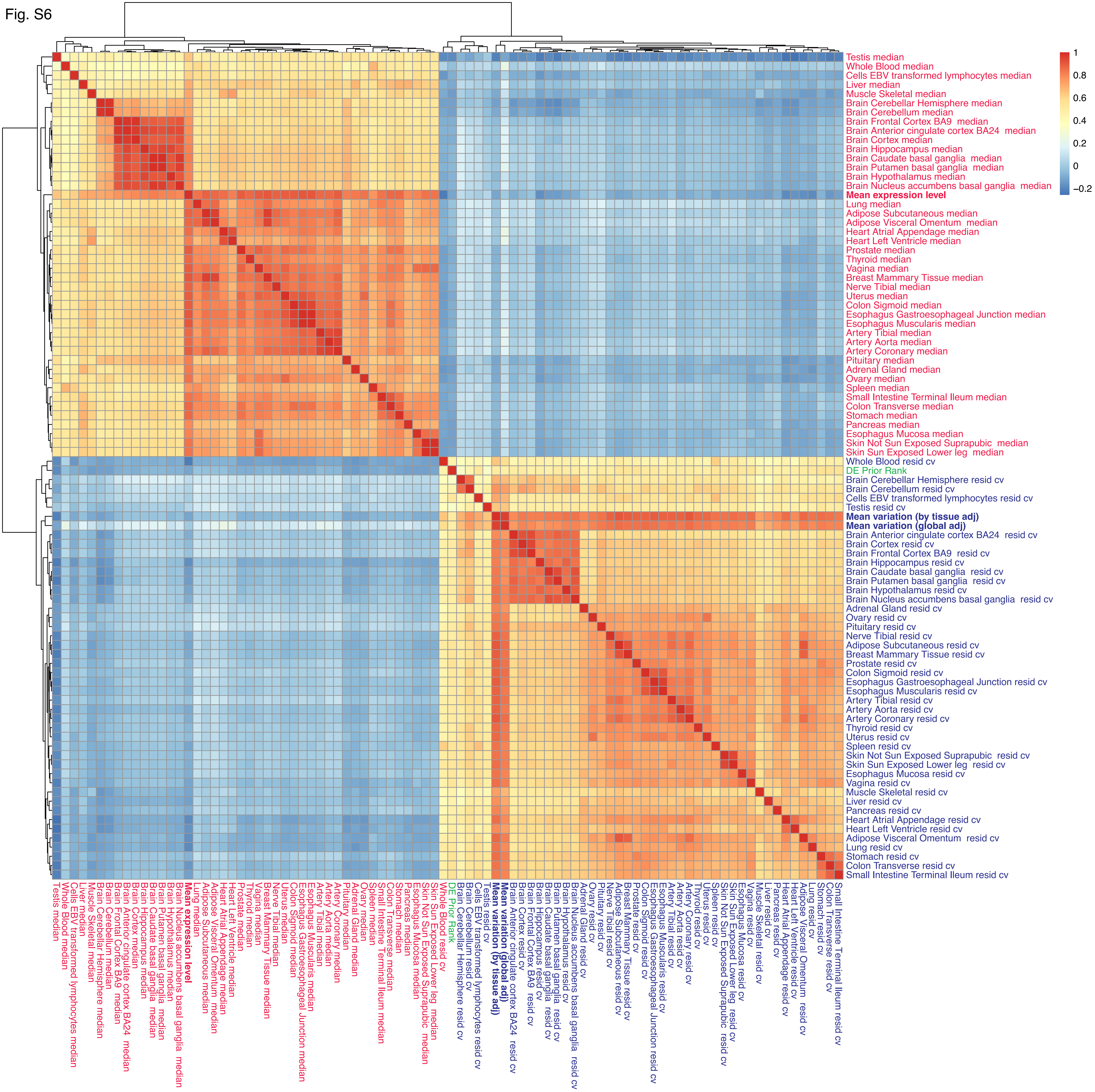
1
0.8
0.6
0.4
0.2
0
−0.2
Testis median
Whole Blood median
Cells EBV transformed lymphocytes median
Liver median
Muscle Skeletal median
Brain Cerebellar Hemisphere median
Brain Cerebellum median
Brain Frontal Cortex BA9 median
Brain Anterior cingulate cortex BA24 median
Brain Cortex median
Brain Hippocampus median
Brain Caudate basal ganglia median
Brain Putamen basal ganglia median
Brain Hypothalamus median
Brain Nucleus accumbens basal ganglia median
Mean expression level
Lung median
Adipose Subcutaneous median
Adipose Visceral Omentum median
Heart Atrial Appendage median
Heart Left Ventricle median
Prostate median
Thyroid median
Vagina median
Breast Mammary Tissue median
Nerve Tibial median
Uterus median
Colon Sigmoid median
Esophagus Gastroesophageal Junction median
Esophagus Muscularis median
Artery Tibial median
Artery Aorta median
Artery Coronary median
Pituitary median
Adrenal Gland median
Ovary median
Spleen median
Small Intestine Terminal Ileum median
Colon Transverse median
Stomach median
Pancreas median
Esophagus Mucosa median
Skin Not Sun Exposed Suprapubic median
Skin Sun Exposed Lower leg median
Whole Blood resid cv
DE Prior Rank
Brain Cerebellar Hemisphere resid cv
Brain Cerebellum resid cv
Cells EBV transformed lymphocytes resid cv
Testis resid cv
Mean variation (by tissue adj)
Mean variation (global adj)
Brain Anterior cingulate cortex BA24 resid cv
Brain Cortex resid cv
Brain Frontal Cortex BA9 resid cv
Brain Hippocampus resid cv
Brain Caudate basal ganglia resid cv
Brain Putamen basal ganglia resid cv
Brain Hypothalamus resid cv
Brain Nucleus accumbens basal ganglia resid cv
Adrenal Gland resid cv
Ovary resid cv
Pituitary resid cv
Nerve Tibial resid cv
Adipose Subcutaneous resid cv
Breast Mammary Tissue resid cv
Prostate resid cv
Colon Sigmoid resid cv
Esophagus Gastroesophageal Junction resid cv
Esophagus Muscularis resid cv
Artery Tibial resid cv
Artery Aorta resid cv
Artery Coronary resid cv
Thyroid resid cv
Uterus resid cv
Spleen resid cv
Skin Not Sun Exposed Suprapubic resid cv
Skin Sun Exposed Lower leg resid cv
Esophagus Mucosa resid cv
Vagina resid cv
Muscle Skeletal resid cv
Liver resid cv
Pancreas resid cv
Heart Atrial Appendage resid cv
Heart Left Ventricle resid cv
Adipose Visceral Omentum resid cv
Lung resid cv
Stomach resid cv
Colon Transverse resid cv
Small Intestine Terminal Ileum resid cv

**Supplementary Figure 6**

Heatmap showing correlations (Spearman correlation coefficient) between expression levels and expression variations across different human tissues as well as their correlation with DE prior from (Crow et al. 2019). Labels contain tissue names from GTEx project; ‘resid cv’ stands for expression variation, ‘median’ indicates median expression level (log-transformed). ‘Mean variation (by tissue adj)’ is mean of expression variation across all tissues where a gene is expressed (corrected for expression level in each tissue separately, Methods). ‘Mean variation (global adj)’ is mean of coefficients of variations in all tissues where a gene was expressed, which was then corrected for mean expression level across the corresponding tissues (Methods). Color code for the labels: red – expression levels; blue – expression variations; green – DE prior. Mean expression level and variation labels are highlighted with bold font.

Fig. S7

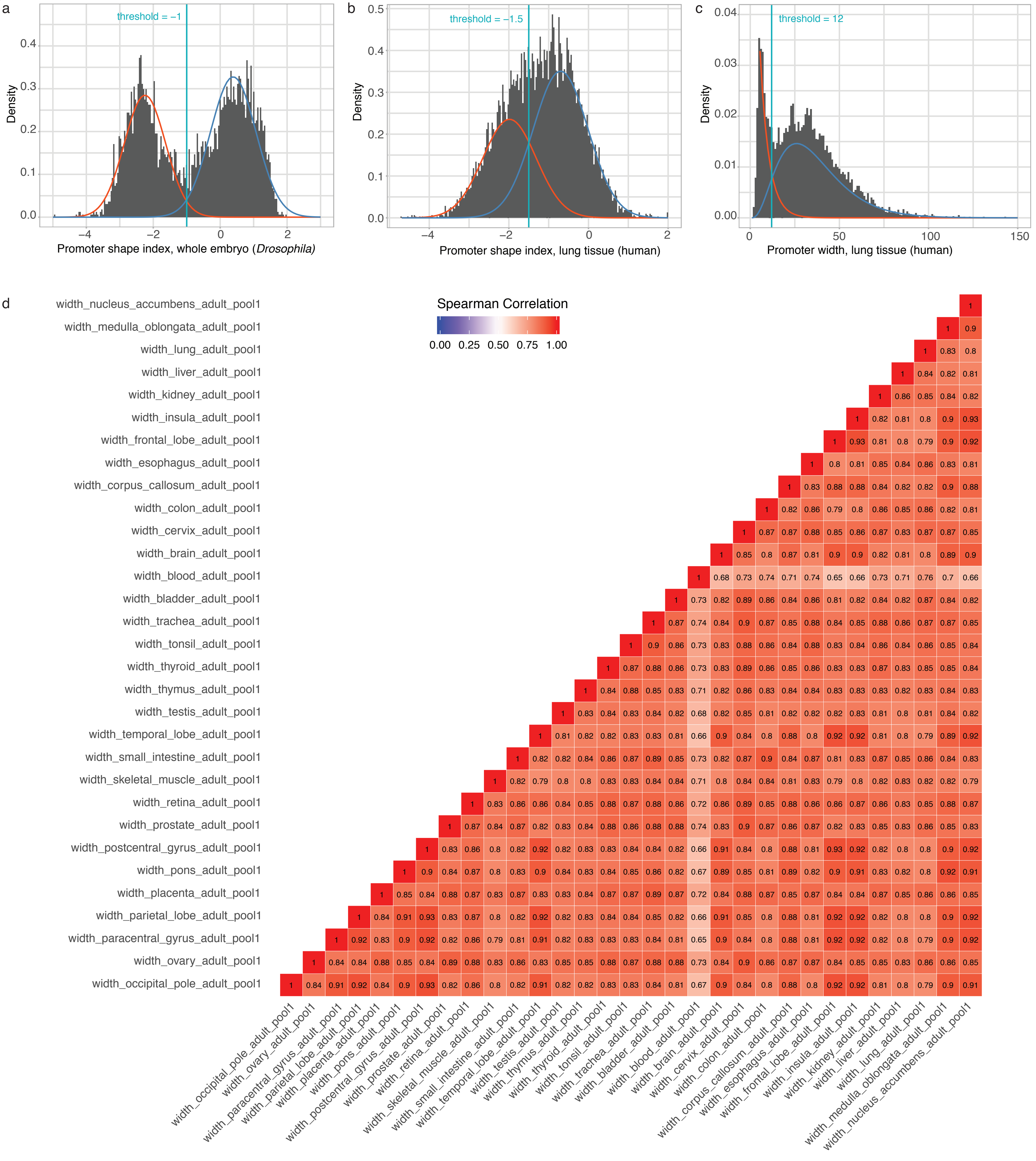
a
threshold = −1
Density
Promoter shape index, whole embryo (Drosophila)
b
threshold = −1.5
Density
Promoter shape index, lung tissue (human)
c
threshold = 12
Density
Promoter width, lung tissue (human)
d
Spearman Correlation
0.00 0.25 0.50 0.75 1.00
width_nucleus_accumbens_adult_pool1
width_medulla_oblongata_adult_pool1
width_lung_adult_pool1
width_liver_adult_pool1
width_kidney_adult_pool1
width_insula_adult_pool1
width_frontal_lobe_adult_pool1
width_esophagus_adult_pool1
width_corpus_callosum_adult_pool1
width_colon_adult_pool1
width_cervix_adult_pool1
width_brain_adult_pool1
width_blood_adult_pool1
width_bladder_adult_pool1
width_trachea_adult_pool1
width_tonsil_adult_pool1
width_thyroid_adult_pool1
width_thymus_adult_pool1
width_testis_adult_pool1
width_temporal_lobe_adult_pool1
width_small_intestine_adult_pool1
width_skeletal_muscle_adult_pool1
width_retina_adult_pool1
width_prostate_adult_pool1
width_postcentral_gyrus_adult_pool1
width_pons_adult_pool1
width_placenta_adult_pool1
width_parietal_lobe_adult_pool1
width_paracentral_gyrus_adult_pool1
width_ovary_adult_pool1
width_occipital_pole_adult_pool1

**Supplementary Figure 7**

**(a-c)** Distribution of promoter shape index and width by gene. **(a)** Histogram of genes' promoter shape index in *Drosophila* embryos (data from (Schor et al. 2017)). **(b)** Histogram of genes' promoter shape index in human lung tissue (Fantom5). **(c)** Histogram of genes' promoter width in human lung tissue (Fantom5). Density lines in **a-c** show fit of mixture distributions, vertical line indicates threshold for separating broad and narrow promoters (Methods). **(d)** Heatmap showing correlations (Spearman correlation coefficient) between promoter widths in different human tissues. Label names contain tissue names according to Fantom5 project.

Fig. S8

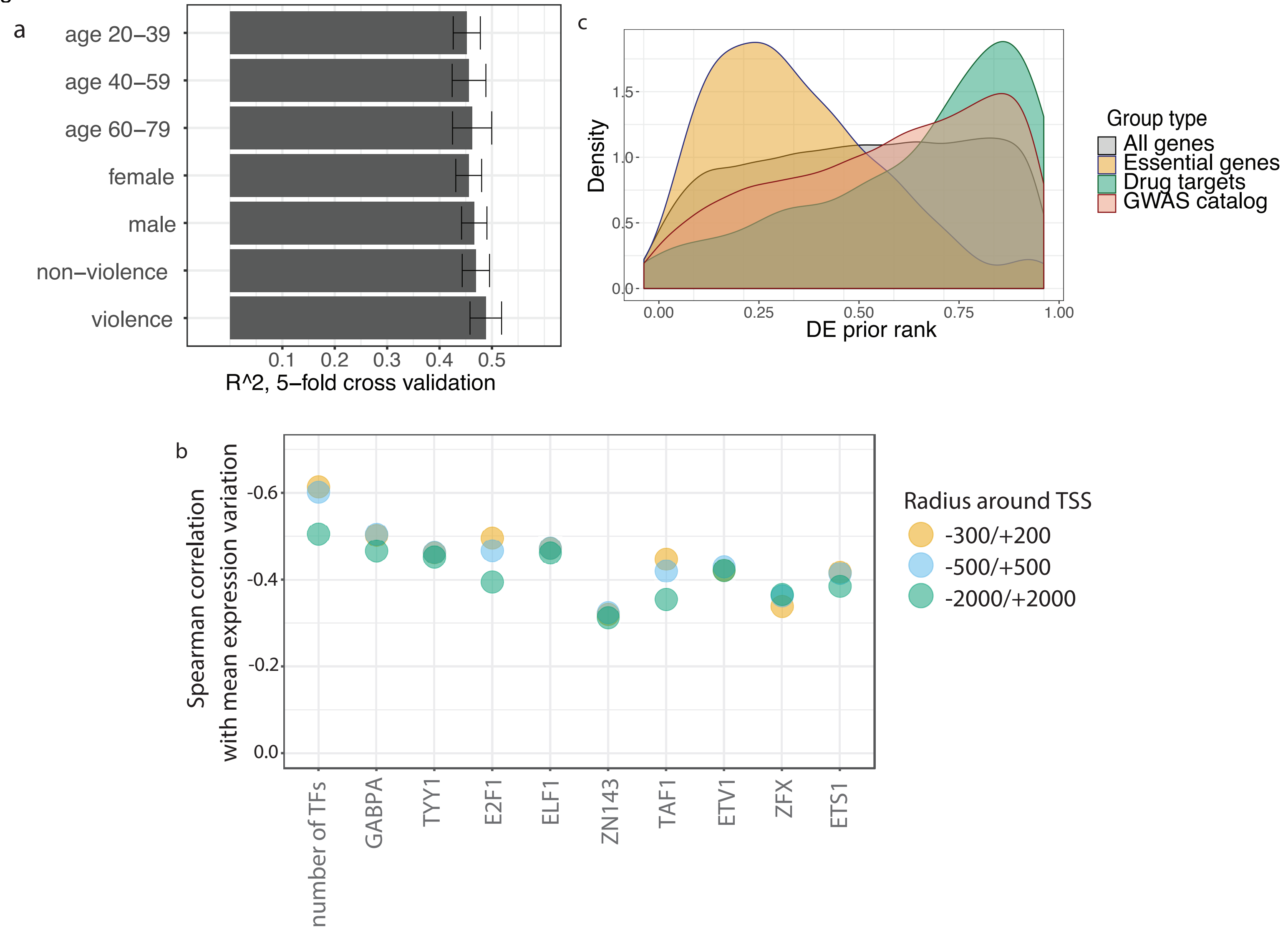

a
age 20–39
age 40–59
age 60–79
female
male
non–violence
violence
0.1
0.2
0.3
0.4
0.5
R^2, 5–fold cross validation
c
Density
1.5
1.0
0.5
0.0
0.00
0.25
0.50
0.75
1.00
DE prior rank
Group type
All genes
Essential genes
Drug targets
GWAS catalog
b
Spearman correlation
with mean expression variation
-0.6
-0.4
-0.2
0.0
number of TFs
GABPA
TYY1
E2F1
ELF1
ZN143
TAF1
ETV1
ZFX
ETS1
Radius around TSS
-300/+200
-500/+500
-2000/+2000

**Supplementary Figure 8**

**(a)** Random forest performance (R^2) for predicting expression variation in lung dataset using different subsets of samples (using samples metadata from GTEx website). Whiskers stand for standard deviation of R^2 obtained from 5-fold cross-validation. **(b)** Spearman correlation coefficient between mean expression variation and TF features dependent on the width of the TSS-proximal region used to associate TFs with genes. Three intervals were considered: -/+500 bp around TSS (used in the main analysis for TFs and chromatin states), -300/+200 (used for some core promoter features, e.g. promoter shape and TATA-box), and -/+2 kB. Correlations are reposted for total number of TFs in the TSS-proximal region and top-10 important TFs for predicting mean expression variation in the main model (based on Boruta feature selection). **(c)** DE prior of specific genes groups (GWAS hits, essential genes, drug targets) compared to the distribution of DE prior for all genes in the dataset.

#### Supplementary Table legends:

**Supplementary table 1.** Features used to predict expression level and variation in *Drosophila*. Columns:
Feature name – feature name as used in the master table (Supplementary tables 2 and 3)
Full name – full feature name
Class – feature class (see Table 1)

**Supplementary table 2.** Full master table including gene symbol and Flybase ID v6.13 (gene_name and gene_id), expression data at three time-points (time) and full feature table (before feature selection and removing several unused features). For each time-point, genes' median expression level (median), final measure of expression variation and expression variation calculated at several intermediate filtering steps (adjustment for expression level dependence performed on the set of genes passing the corresponding filtering step, Methods):

1. resid_cv_extrRm – set of genes after removing top and bottom-5% by expression level;
2. resid_cv_decrExprFilt - set of genes from 1) after removing genes that decreased in expression between 10-12 and 2-4 hours;
3. resid_cv_decrExprFilt_filtNA - set of genes from 2) after removing genes with missing values in the feature table (final variation measure)

Feature names used in the random forest are explained in Supplementary table 1.
Several additional features that were excluded from the random forest are provided gene: gene start and end coordinates (start, end), ID of topologically associated domain from several TAD annotations (tad_id.ramirez and tad_id.2_4h, Methods), expression log2-fold change between different time-points (expr_l2fc_10vs2h, expr_l2fc_6vs2h, expr_l2fc_10vs6h)

**Supplementary table 3.** Final feature table including gene symbol and Flybase ID v6.13 (gene_name and gene_id), all features (before feature selection), median expression level (median), expression coefficient of variation (cv), expression variation (resid_cv, same as resid_cv_decrExprFilt_filtNA in Supplementary table 2) at 10-12h. Feature names are explained in Supplementary table 1.

**Supplementary table 4.** Feature importance scores (from Boruta) and correlations with predicted variables (Methods). Only features important in at least one prediction are included. NAs indicate non-significant features in the corresponding predictions.
Columns 1-3 are the same as in Supplementary Table 1. Columns 4-18:
med_imp_var - median feature importance for predicting expression variation
med_imp_med - median feature importance for predicting median expression level
med_imp_shape_ind - median feature importance for predicting promoter shape index
med_imp_broad_var –- median feature importance for predicting expression variation in broad promoter genes
med_imp_narrow_var - median feature importance for predicting expression variation in narrow promoter genes
med_imp_narrow_lev - median feature importance for predicting median expression level in narrow promoter genes
med_imp_broad_lev - median feature importance for predicting median expression level in broad promoter genes

cor_var - feature correlation with expression variation
cor_med - feature correlation with median expression level
cor_shape_ind - feature correlation with promoter shape index
cor_var_broad - feature correlation with expression variation in broad promoter genes
cor_var_narrow - feature correlation with expression variation in narrow promoter genes
cor_med_broad - feature correlation with feature correlation with expression variation in broad promoter genes
cor_med_narrow - feature correlation with feature correlation with expression variation in narrow promoter genes

**Supplementary table 5.** Results of Fisher's exact test for feature enrichments in specific groups of genes. Gene groups are defined in Fig. 3a. Feature names are explained in Supplementary table 1. Columns 2-6 contain the values from the contingency tables used in the Fisher's test. Odds ratios from the table are visualized in Fig. 4c. Columns:
comp – groups of genes being compared, e.g. broad promoter genes vs. narrow promoter genes (broad_vs_other) or 'narrow-low' genes vs. 'narrow-high' and 'broad' genes (narrow-low_vs_other).
Feature – feature for which the enrichment was calculated, e.g. presence of peaks for MESR4 in the TSS-proximal regions (modERN.MESR4.E0_24.prox)
num_group_feature – number of genes in the tested group having the feature
num_nongroup_feature - number of genes in the other group having the feature
num_group_nofeature - number of genes in the tested group not having the feature
num_nongroup_nofeature - number of genes in the other group not having the feature
pval - the p-value of the test
odds_ratio - an estimate of the odds ratio using conditional Maximum Likelihood Estimate (fisher.test function, R package *stats*)
pval_bh - the p-value after correction for multiple testing (Benjamini-Hochberg correction)
num_comp – number of comparisons, used in Benjamini-Hochberg correction. Enrichments of TFs (modERN.*tf_name*.prox) and gene categories were tested separately, hence number of comparisons differs.

**Supplementary table 6.** Gene Ontology (GO) functional enrichment of genes grouped into eight clusters by promoter shape (broad/narrow) and expression variation (four quantiles by expression variation calculated separately within broad and narrow promoter genes). Exact values for quantiles are provided in Methods. Results are provided from compareCluster function (R package *clusterProfiler*). Columns:
Cluster - gene group tested for enrichment, e.g. 1.broad (bottom-25% by expression variation within broad promoter genes) or 4.narrow (top-25% by expression variation within broad narrow genes)
ID - GO category ID
Description – GO category description
GeneRatio – gene ratio
BgRatio – background ratio
pvalue - enrichment p-value
p.adjust – adjusted p-value (Benjamini-Hochberg correction)
qvalue - enrichment q-value
geneID - entrez IDs of genes from the GO category in the tested group
Count - number of genes from the GO category in the tested group
Ontology – Biological Process (BP) or Molecular Function (MF)

**Supplementary table 7.** GO functional enrichment of genes grouped into four clusters by expression log2-fold change between 10-12h and 2-4h after fertilization (intervals are provided in column *Cluster*). Genes with log2-fold change below 0 were excluded from final analysis (Methods). Column names are similar to Supplementary table 6.

**Supplementary table 8.** List of GTEx tissues used in the analysis (RNA-seq data).

**Supplementary table 9.** Tissue-specific expression data, DE prior (DE_Prior_Rank) and several gene annotations (essential genes, drug targets, GWAS hits; for details see Methods). Columns names for tissue-specific expression contain tissue names as listed in Supplementary table 8. NA indicates that a gene is not expressed (or did not pass filtering criteria) in the corresponding tissue. Columns:
Gene_name – gene symbol;
Gene_id – Ensembl gene ID;
*Tissue_name*_mean – gene mean expression level (log-transformed) across individuals in the corresponding tissue;
*Tissue_name*_median - gene median expression level (log-transformed) across individuals
*Tissue_name*_cv - gene coefficient of variation across individuals
*Tissue_name*_recid_cv - gene expression variation across individuals (final measure of variation, adjusted for median dependence);
DE_Prior_Rank – Differential expression prior from (Crow et al. 2019);
GWAS_Upstream_gene_id and GWAS_Downstream_gene_id – EBI GWAS catalog genes (upstream or downstream of GWAS hits);
CEGv2_subset – essential genes;
Drug_targets_nelson and FDA_approved_drug_targets – drug targets.

**Supplementary table 10.** Features used to predict expression level and variation in human. Columns:
Feature name – feature name as used in the master table (Supplementary tables 2 and 3)
Feature class – e.g. transcription factors or chromatin states
Feature type – tissue-specific (only used in Supplementary tables 13-15), non-tissue-specific or averaged across tissues (Methods)

**Supplementary table 11.** List of Pantom5 tissues used in the analysis (CAGE data).

**Supplementary table 12.** List of tissues with chromatin states used in the analysis (chromHMM)

**Supplementary table 13.** Feature table for predicting expression variation (resid_cv) and median expression level (median, log-scale) across individuals in Lung tissue. Features used to predict expression variation and level are listed in Supplementary table 10.

**Supplementary table 14.** Feature table for predicting expression variation (resid_cv) and median expression level (median, log-scale) across individuals in Muscle tissue. Features used to predict expression variation and level are listed in Supplementary table 10.

**Supplementary table 15.** Feature table for predicting expression variation (resid_cv) and median expression level (median, log-scale) across individuals in Ovary tissue. Features used to predict expression variation and level are listed in Supplementary table 10.

**Supplementary table 16.** Feature table for predicting mean expression variation (mean_variation) and mean expression level (mean_median, log-scale) aggregated across all expressing tissues (Methods). Features used to predict expression variation and level are listed in Supplementary table 10.

**Supplementary table 17.** Feature importance scores (from Boruta) and correlations with predicted variables (Methods). Only features important in at least one prediction are included. NAs indicate non-significant features in the corresponding predictions.
Columns 1-3 are the same as in Supplementary Table 1. Columns 4-18:
